# Resolving spatial complexities of hybridization in the context of the gray zone of speciation in North American ratsnakes (*Pantherophis obsoletus* complex)

**DOI:** 10.1101/2020.05.05.079467

**Authors:** Frank T. Burbrink, Marcelo Gehara, Edward A. Myers

## Abstract

Inferring the history of divergence between species in a framework that permits the presence of gene flow has been crucial for characterizing the gray zone of speciation, which is the period of time where lineages have diverged but have not yet achieved strict reproductive isolation. However, estimates of both divergence times and rates gene flow often ignore spatial information, for example the formation and shape of hybrid zones. Using population genomic data from the eastern ratsnake complex (*Pantherophis obsoletus*), we infer phylogeographic groups, gene flow, changes in demography, the timing of divergence, and hybrid zone widths. We examine the spatial context of diversification by linking migration and timing of divergence to the location and widths of hybrid zones. Artificial neural network approaches are applied to understand how landscape features and past climate have influenced population genetic structure among these lineages prior to hybridization. Rates of migration between lineages are associated with the width and shape of hybrid zones. Timing of divergence is not related to migration rate across species pairs and is therefore a poor proxy for inferring position in the gray zone. However, timing of divergence is related to the number of loci weakly introgressing through hybrid zones.

---

Wide-ranging species complexes that cross numerous biogeographic barriers provide opportunities to better understand how changing landscapes affect diversification, gene flow, demography, and the formation of hybrid zones. Lineages within species complexes ranging across heterogeneous landscapes are likely at different stages of the speciation process. These differences may reflect how specific environmental and biogeographic barriers have uniquely altered changes in gene flow and other demographic processes in a particular complex (Myers et al. 2020). Even at the same barrier within related groups of organisms, the timing of divergence and degree of gene flow can be species specific and dependent on when they encountered the barrier (Riddle 2016; Myers et al. 2019b). Alternatively, generalist species not as constrained to specific habitats may not show a correlation between genetic variation and landscape features (Joseph and Wilke 2007; Makowsky et al. 2009; Lourenço et al. 2017).

Lineage divergence can be placed in the context of the gray zone of speciation, which defines the range of time where speciation proceeds from early population differentiation with unfettered gene flow to nearly reproductive isolation defined by low rates of gene flow (de Queiroz 2007; Hewitt 2008; Roux et al. 2016; Jackson et al. 2017). The ability to delimit species therefore may be correlated with their position in the gray zone, which has previously been assessed by relating measures of genetic isolation and migration (the genealogical divergence index; GDI) with species-delimitation probabilities (Roux et al. 2016; Leaché et al. 2019).

Moreover, the position within the gray zone could be simply predicted by the age of lineage formation relative to population size if the evolution of Dobzhansky-Muller incompatibilities via genetic drift alone occurred after divergence (Orr 1995; Gavrilets 2004; Singhal and Moritz 2013). Deviations from a negative correlation between rates of gene flow and age of divergence could occur where increased gene flow between the oldest divergences in a group may cause lineages to rapidly collapse, such as in the case where physical barriers to gene flow disappear. On the other hand, strong selection may prevent young lineages from exchanging alleles at particular loci, shortening the time in the gray zone (Mayr 1963; Barton 2010; Feder et al. 2012; Roux et al. 2016; Edwards et al. 2020). Therefore, finding a threshold that can determine if two species are unique given only divergence times may fail where rates of gene flow between lineages drastically change over time and space (Gourbière and Mallet 2010; Nosil et al. 2017).

Understanding how divergence has occurred over the landscape in the context of hybrid zones should provide a better understanding of the location of diverging lineages in the gray zone. As first point of inquiry, examining gene flow in space determines if lineages were generated via isolation and migration or are simply artificially partitioned groups with continuous gene flow over the landscape defined by isolation by distance (IBD; Wright 1943; Frantz et al. 2010; Bradburd et al. 2018). Barring the latter, if species pairs rapidly adapt to unique habitats across their distribution, then age of divergence and degree of gene flow may be unrelated relative to neutral processes when compared across taxa (Nosil and Crespi 2006; Agrawal et al. 2011; Nosil 2012; Karrenberg et al. 2019). Therefore, divergence over landscape can show how selection at a barrier can be discordant from the timing of origin of the lineages (Barton and Hewitt 1985; Harrison 1993; Jiggins and Mallet 2000; Gay et al. 2008; Seehausen et al. 2014; Stankowski et al. 2017).

To understand population divergence many studies use coalescent-derived, isolation-migration models (Hey and Nielsen 2004; Hey 2010) to estimate gene flow throughout time, reflecting position in the gray zone of speciation. These coalescent models have been used to detect unique species (Rannala and Yang 2020), however, they lack any spatial component necessary to understand the underlying geographic and environmental processes influencing contemporary migration and degree of reproductive isolation reflected within any particular locus. Coalescent estimates for understanding gene flow, timing, and species delimitation are also affected by geographic sampling and connectivity among populations (Pante et al. 2015; Coates et al. 2018; Mason et al. 2020). Studying this connectivity can determine if hybridization among lineages are enhancing or reducing reproductive isolation (Abbott et al. 2013) and can reveal if contemporary gene flow is restricted to a few loci (genic species) or most of the genome (biological species; Nosil and Feder 2013). Therefore, studying hybrid zones should enhance traditional coalescent estimators of divergence by determining: 1) the location and size where lineages meet and migration occurs, 2) how genes respond to selection across these zones, 3) the degree of contemporary reproductive isolation, and 4) if parental lineages exist in unique habitats.

To understand the factors shaping reproductive isolation in a species complex, we ask whether divergence time predicts rates of gene flow or cline width across space in the North American ratsnakes (*Pantherophis obsoletus* complex). This complex is a wide-ranging group of four species found throughout the forested regions of the Eastern Nearctic and nearby Chihuahuan Desert that have diverged at three unique biogeographic barriers. The primary divergence in this complex occurred at the Mississippi River, which has consistently been identified as one of the main biogeographic barriers in the ENA (Robison 1986; Burbrink et al. 2000; Soltis et al. 2006; Brandley et al. 2010; Zellmer et al. 2012; Myers et al. 2020). Further east, the Appalachian Mountains and Apalachicola/Chattahoochee River System (AARS) have noted barrier effects in other organisms and likely contributed to the divergence between *P. alleghaniensis* and *P. spiloides* (Walker and Avise 1998; Burbrink et al. 2000; Soltis et al. 2006). West of the Mississippi River, divergences occurred at the transition between temperate forests to the western edge of the Edwards Plateau into the Chihuahuan Desert and isolated the species *P. obsoletus* and *P. bairdi* (Fig.1; Lawson and Lieb 1990; Burbrink et al. 2000; Burbrink 2001).

**Fig 1.**
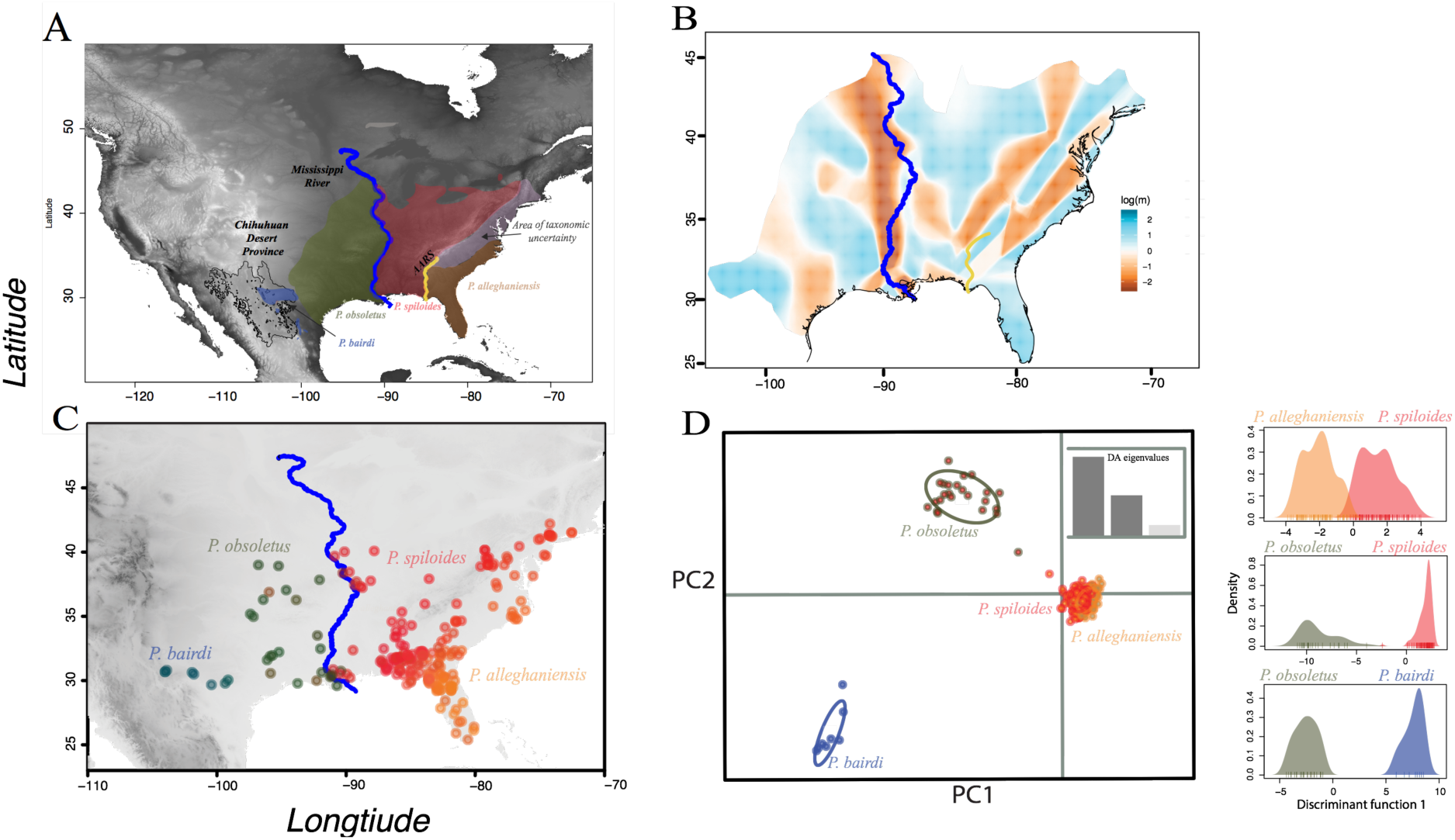
Distribution of species within the North American ratsnake complex. (A) Map showing range and geographic features: Chihuahuan Desert Province, the Mississippi River, and the Appalachian Mountains and Apalachicola/Chattahoochee River System (AARS). (B) Estimated effective migration surfaces (EEMS) showing areas of both high (blue) and low migration (brown). (C) Population structure using Discriminant Analysis of Principal Components (DAPC) shows the location of four geographically distinct lineages. (D) Bi-plots of PC space shows relative distances among the four taxa using DAPC.

Since the Pliocene, the ENA has experienced numerous climate change events associated with glacial cycles that forced species into refugia and compressed populations; upon climate amelioration populations expanded into formerly glaciated areas (Hewitt 2000; Bintanja and van de Wal 2008; Burbrink et al. 2016). Using dense population sampling and genomic-scale data, we combine isolation-migration estimates with spatial information to understand species divergence in the context of geographic distance, contemporary and historical environments, and biogeographic boundaries. We examine how migration rates and divergence dates relate to the widths of hybrid zones at biogeographic boundaries and investigate demographic changes through time to understand if population expansion occurs among all lineages as predicted given glacial cycling. Addressing these questions provides an integrated view of phylogeographic history over a physically and historically complex landscape and yields a clearer understanding of speciation processes and delimitation in the gray zone.

## Methods

### Dataset

We sampled 288 individuals liberally covering the range of all species within the *Pantherophis obsoletus* complex (Fig.1; Dryad XXX). DNA was extracted from all samples using Qiagen DNeasy Blood & Tissue Kits and samples were screened for quality using broad-range Qubit Assays. We used services from RAPiD Genomics (https://www.rapid-genomics.com/services/) to generate 5472 baits and to sequence 5060 conserved elements (UCEs) loci following the protocols from (Faircloth et al. 2012) and (Sun et al. 2014). These markers have been used to address phylogeographic/population genetic and deeper phylogenetic questions (Myers et al. 2019a; Younger et al. 2019). We mapped UCE reads to a Chromium 10x *Pantherophis spiloides* genome (in prep), removed loci containing >50% missing samples, and removed individual specimens missing >30% of all alignments (details on the assembly of UCE loci are available in Supporting Information Material 1). We produced both phased locus datasets, used in our coalescent estimators, and single SNPs/locus for all other analyses. We filtered the unlinked SNP dataset to remove alleles with a minor allele frequency <0.1 and used these data for population structuring analyses (Linck and Battey 2019).

### Geographic groupings

We estimated population structure by comparing Discriminant Analysis of Principal Components (DAPC; Jombart et al., 2010) in the R package (R Core Team 2015) adegenet v2.1.2 (Jombart 2008). Additionally, we estimated effective migration surfaces (EEMS; Petkova et al., 2016) three times with different starting seeds to insure consistency. This method models effective migration rates over geography to represent regions where migration is low in cases where genetic dissimilarity increases rapidly (details on these population grouping methods and other methods used for assessing structure are available in Supporting Information Material 1).

### Isolation, migration, and historical demography

We generated a coalescent-based species tree with SNAPP v1.3.0 (Bryant et al. 2012) to test species limits and understand relationships among the four groups identified from clustering analyses. Because generating a tree in SNAPP using all individuals was computationally intractable, we used four individuals per taxon that were sampled from different regions of the taxon’s distribution. We estimated SNAPP trees using BEAST v2.5.2 (Bouckaert et al. 2014; details on parameter setting for SNAPP are available in Supporting Information Material 1). We assessed four alternative species delimitation models including four taxa, three taxa (collapsing *P. bairdi/P. obsoletus* or *P. alleghaniensis/P. spiloides*), and two taxa (collapsing *P. bairdi/P. obsoletus* and *P.alleghaniensis/P. spiloides)*.

Using the four inferred lineages, we examined which demographic models of species divergence best described the origins of these groups given divergence time, historical demographic change, and migration between spatially adjacent taxa. We tested 2,300 candidate isolation-migration models including all possible topologies with or without migration between all spatially adjacent pairs using the program PHRAPL (Jackson et al. 2017) implemented in R. To further test that four genetic lineages exist, we ran pairwise comparisons between the best selected model and models of three populations where sister species were collapsed into a single entity; four *Pantherophis* species against one model where *P. obsoletus* and *P. bairdi* were collapsed, and another where *P. spiloides* and *P. alleghaniensis* were collapsed (details on PHRAPL runs are available in Supporting Information Material 1).

To estimate gene flow, timing of divergence, and demographic change, we used PipeMaster (Gehara et al., 2017; Gehara et al. in review) to simulate genetic data and perform approximate Bayesian computation (ABC) and supervised machine-learning. Here we used the best model selected in the PHRAPL analysis as a template (same topology and migration parameters) to generate three competing models: (i) an isolation migration model with constant population size for each lineage and constant migration, (ii) demographic change along each lineage and constant migration through time and demographic change, and (iii) population size change with migration occurring after the Last Glacial Maximum. We simulated 100,000 data sets of 54 summary statistics and performed ABC rejection with 0.01 tolerance level to select the best of these three models (Table S1). We then took the selected model and performed rejection using a 0.1 tolerance level with a neural network regression using a single layer and 20 nodes to estimate the model parameters. We also performed this without hybrid individuals (admixture proportions between 0.5-0.7) to explore how including these samples may change estimates of divergence timing, demographic change, timing of demographic change, and migration within these lineages.

**Table 1.** Measures of divergence dates in millions of years (My), migration rates (first number is first taxon migrating into second taxon, e.g., *Pantherophis alleganiensis* into *P.spiloides*, and second represents second species migrating into the first), and cline widths for both spatial distance (Km) and environmental distance for all species-pair comparisons in the *Pantherophis obsoletus* complex. Percentage of median width over total sampled distance are shown for both spatial and environmental clines

| Group | Age (My) | Migration Rate (ind/gen) | Cline Width (Km) | Cline Width (Env) |
| --- | --- | --- | --- | --- |
| Eastern (*P. alleghaniensis* x *P.spiloides*) | 1.95 (0.674-0.2.95) | 2.79/3.79 | 256 (168-356); 14.8% | 0.81 (0.32-1.45); 4.0% |
| Central (*P.spiloides* x *P.obsoletus*) | 3.18 (1.65-5.02) | 1.96/1.28 | 147 (40-283); 7.2% | 0.13 (0.06-0.24); 5.0% |
| Western (*P.obsoletus* x *P. bairdi*) | 1.10 (0.407-02.62) | 0.65/0.55 | 21 (2-46); 1.6% | 4.4 (3.2-12.53); 2.1% |

### Spatial population genetics

We used redundancy analyses (RDA; Legendre et al. 2011; Diniz-Filho et al. 2013) and artificial neural networks (ANN; Lek and Guégan 1999; Legendre and Fortin 2010; Legendre et al. 2011) to understand how contemporary and historical aspects of the landscape and climate generated genetic diversity. To set up our data for these analyses, we used as our response variable Euclidian genetic distance and transformations of this (see below) among all individuals in R. For predictor variables, we used 1) geographic distances measured as pairwise great-circle distances among all points to account for isolation-by-distance using the R package fossil (Vavrek 2020), 2) binary categorization of isolation east and west of the Mississippi River, 3) elevation, and 4) bioclimatic variables at 2.5 mins representing current climate, Last Glacial Maximum (LGM, 21kya), Last interglacial (LIG, 130 kya), and Pleistocene Marine Isotope Stage 19 (MS19, 787 kya) all obtained from the PaleoClim database (Brown et al. 2018). We also extracted level II ecoregions for all individuals. Ecoregion variables delineate unique ecological areas and are well characterized across North America (Bailey 1995). For the bioclimatic data and elevation, we extracted parameter values given the latitude and longitude for each individual sample using the R package raster (Hijmans et al. 2014). To reduce dimensionality of the bioclimatic variables for each type (current, LGM, etc.), we eliminated correlated variables (>0.90) and transformed remaining variables using centered principal components analyses in R. We transformed categorical ecoregion data into dummy variables and reduced dimensionality using logistic PCA for binary variables (Landgraf and Lee 2015). To account for the effect of geographic distances on genetic structure, we transformed spatial distance between all samples using principal coordinates neighbor matrices (PCNM) in the R package vegan (Dixon 2003). This method allowed us to account for multiple spatial structures (neighborhoods) on heterogenous ecological structures and barriers, which better clarified non-spatial effects on genetic structure (Borcard and Legendre 2002).

We first used RDA, asymmetric canonical analysis, to determine the significance for each spatial predictor axis (Legendre et al. 2011) on genetic structure (McGaughran et al. 2014; Noguerales et al. 2016). We then used the reduced set of significant spatial axes to examine their effect with environmental and barrier variables on genetic structure using RDA and ANN (see Supporting Information Material 1 for details). Machine learning methods, such ANN, can infer non-linear interactions among many predictor variables, regardless of the distribution or variable type and can generate a range of models of varying complexity (Lek et al. 1996; Zhang 2010; Libbrecht and Noble 2015; Sheehan et al. 2016). We transformed Euclidian genetic distances into principal coordinates (PCoA; (Gower 1966) in adegenet (Jombart 2008) to produce uncorrelated orthogonal axes to summarize genetic variation and assessed accuracy and variable importance on the first 10 PCoA axes. We used regression-based ANN to determine which variables predict genetic distances and rank those with the highest model importance. To conduct the ANN regressions we used the R package *caret* (Kuhn 2008) and partitioned the data into the standard 70% training and 30% test sets (Lek et al. 1996; Zhang 2010; Burbrink et al. 2020).

This analysis was run using 1,000 maximum iterations to ensure convergence. We resampled the data using the default 25 bootstrap replicates to reach convergence over the following parameters: weight decay, root mean squared error, *r*^2^, and mean absolute error. Accuracy was compared to an ANN run with random shuffling of the response variable. We also compared results from our ANN analysis to those using redundancy analyses (RDA, see Supporting Information Material 1 for details).

### Hybrid-zone dynamics

Using the program HZAR v.2-9 (Derryberry et al. 2014) we used admixture proportions to determine if cline widths differ among adjacent species. We used TESS3r to estimate admixture for each species pair using SNPs and geographic data (see Supporting Information Material 1; Caye et al. 2018). As required by HZAR, we reduced 2-dimensional space (latitude and longitude) into 1-dimensional distance from the hybrid zone. We first determined the location of the hybrid zone by mapping admixed individuals (0.5-0.6 admixture), calculating all sample distances to the hybrid zone, and then fixing a sign to these distances where 0 was the hybrid zone, positive distances trended towards one parental taxon, and negative distances trended towards the other. Using the Gaussian cline model, we estimated the center and width of the cline and determined if these sigmoidal distributions have significant tails by fitting the following models: 1) no tails, 2) right tail only, 3) left tail only, 4) mirrored tails, and 5) both tails estimated independently (see Derryberry et al. 2014). We fit these models to our data using AICc and ran the MCMC chains for 5x10^6^ generations, thinned by 5x10^3^ generations, and estimated stationarity using ESS >200 in the R package CODA (Plummer et al. 2006).

To examine the width of these clines over environmental space, we transformed environmental space to a single dimension and measured distance relative to the hybrid zone. We filtered correlated bioclim data (>0.90), reduced dimensionality using PCA, estimated absolute environmental distance between all individual points and hybrid zones, and provided a positive or negative sign depending on the side of the hybrid zone. We ran HZAR described above for spatial distance data.

To examine the width of allelic clines through hybrid zones relative to overall admixture, we also estimated cline widths for each locus over geographic and environmental distance. Here we used allele frequencies for each individual and applied the same distances as described above using HZAR. To better understand the effects of selection over geographic space for each species-pair comparison (see Stankowski et al. 2017), we determined which loci were fixed for each parental taxa (>0.80) in the tails of the cline and also showed sharp clines (reduced widths). The difference in the frequency of fixation (ΔP) between each tail of the cline (sampling 5% of each most distant tail) was estimated and widths were compared to neutral expectations of size (see below).

Using equations from Barton and Gale (1993) and applied in Bailey et al. (2015) we evaluated selection against hybridization for each cline. With estimates of the root mean squared (RMS) maximum lifetime dispersal (σ) for ratsnakes, we predicted the width of the cline under neutrality over a range of generation times (*T*) since the origin of each sister pair and group using the equation: 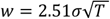. Unfortunately RMS of dispersal is unknown for ratsnakes, but we bound this from 1.02 to 4.03 km given mean and maximum distances of nesting sites to hibernaculum (Blouin-Demers and Weatherhead 2002). The maximum width was likely an underestimate of lifetime dispersal, particularly in areas in the southern United States where ratsnakes are not constrained by hibernacula.

All scripts, genetic, and spatial data are available on Dryad XX.

## Results

### Data and population genetic structure

After filtering loci for presence in 70% or more individuals, we generated 2,491 UCE loci. For all downstream analyses we randomly sampled one SNP per locus yielding a total of 846 SNPs for 238 individuals for an average of 13.96% missing data.

Both EEMS and DAPC found population structure that generally matched the geographic ranges of *P. alleghaniensis*, *P. spiloides, P. obsoletus*, and *P. bairdi* (Fig. 1 and Fig. S1; Burbrink, 2001). All three EEMS generated similar acceptance proportions for all proposal types (12-51%). Total DAPC assignment probabilities without a priori species groupings were 0.975 (*P. bairdi* = 1.0, *P. obsoletus* = 0.96; *P. spiloides* = 1.0, and *P. alleghaniensis* = 0.97). Similarly, all three EEMS replicates showed low migration at the Mississippi River (MR) and the area separating the *P. bairdi* and *P. obsoletus* in west Texas. East of the MR, estimated low migration occurred near the Appalachian Mountains, though this area is a complex mixture of isolation and gene flow (Fig. 1).

These estimates of population structure all showed *P. alleghaniensis* as occurring mostly in the Florida Peninsula and north along the east coast to Virginia/Delaware, *P. spiloides* was found from western Florida to the Mississippi River up through the Midwest and east to the northeastern US, *P. obsoletus* was found west of the Mississippi River in the forested regions of the Midwest, and *P. bairdi* was distributed in the Chihuahuan Desert of west Texas (Fig. 1).

### Isolation-migration processes

Our SNAPP analyses produced a phylogeny with a root separating the taxa east and west of the Mississippi River with a sister relationship between *P. bairdi* and *P. obsoletus* and then *P*. *spiloides* and *P. alleghaniensis* (Fig. 2), consistent with Burbrink et al. (2000). Bayes factors (BF) showed the four-taxon model as decisively superior to the three-taxon (BF =35.40) and two-taxon model (BF=2022.46).

**Fig 2.**
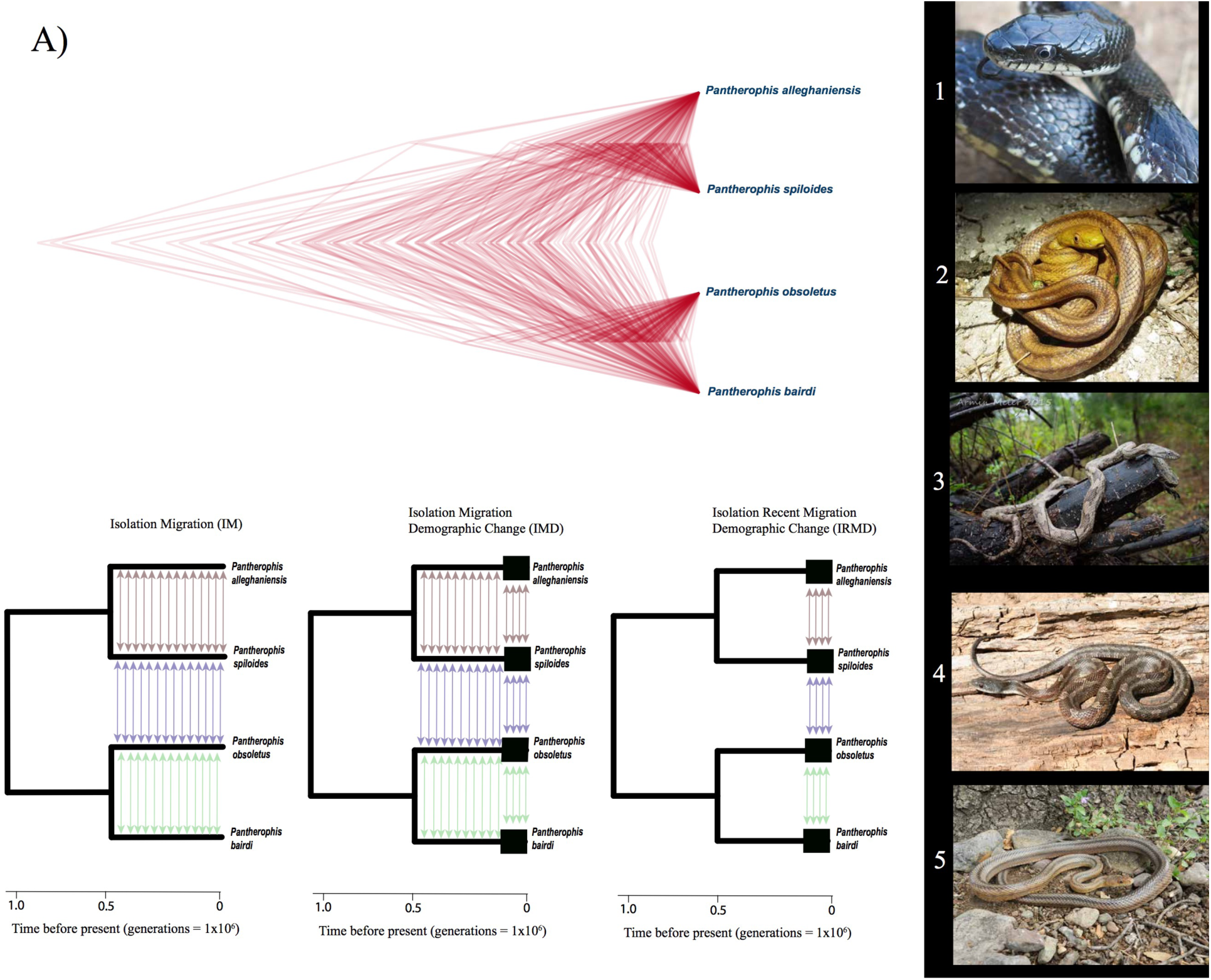
The results of SNAPP species-tree estimation for the four taxa over all loci (A) and models showing Isolation Migration (IM) and Isolation Migration with Demographic Change (IMD; best-fit model) and Isolation with Recent Migration and Demographic Change (IMRD) estimated in PipeMaster. Colored arrows represent migration among spatially adjacent pairs. (B). Photographs representing some of the typical color patterns seen in ratsnakes: 1) *Pantherophis alleghaniensis* (photo F. Burbrink), 2) *P. alleghaniensis* (photo N. Claunch), 3) *P. spiloides* (photo A. Meier), 4) *P. obsoletus* (photo D. Shepard), and 5) *P. bairdi* (E. Myers).

Using those four lineages we filtered models of divergence using PHRAPL. A model delimiting all four taxa, incorporating migration between spatially adjacent lineages was preferred with the same topology as found using SNAPP. ΔAIC between the best ranked model and next model was > 20, suggesting high confidence in model selection (Table S2).

The demographic model estimated from PipeMaster that incorporated historical population size change and constant migration (IMD) best fit the data (posterior probability 1.0; see PCA plots in Fig. S2; Fig. 2). Median migration rates were highest between eastern groups (*P. alleghaniensis* to *P. spiloides* = 2.79 and *P. spiloides* to *P. alleghaniensis* = 3.79 individuals/generation), lower in the central groups across the Mississippi River (*P. spiloides* to *P. obsoletus* = 1.95 and *P. obsoletus* to *P. spiloides* = 1.28 individuals/generation), and lowest in western groups (*P. bairdi* to *P. obsoletus*= 0.55 and *P. obsoletus* to *P. bairdi =* 0.65 individuals/generations; Table 1 & Fig. 3). Divergence occurred earliest in the central lineages at the Mississippi River at 3.181 My (95% quantile =1.65 to 5.02 My), then the eastern lineages at 1.95 My (95% quantile = 0.674 to 2.95 My) and western lineages at 1.10 My (95% quantile 0.407 to 2.62 My; Table 1 & Fig. 3). Divergence dates and average rates of migration were uncorrelated (ρ=0.264, *P*=0.83, n=3). For all taxa we inferred population expansion from ancestral *N_e_* sizes ranging from 5,200 to 5,600 to modern sizes at 138,000 to 281,000 (Fig. 3). Median expansion times were similar for all taxa (0.230-0.260 Ky) with lower 95% quantiles always occurring in the Pleistocene (0.024-0.38 Ky; Fig S3). Posterior distributions for all parameters were different from the uniform priors, though wide distributions of several parameters suggested some uncertainty in these estimates (Fig. 3). Similarly, rerunning PipeMaster without heavily admixed individuals resulted in similar predictions but with lower estimates of migration for eastern and central lineages (Fig. S3).

**Fig 3.**
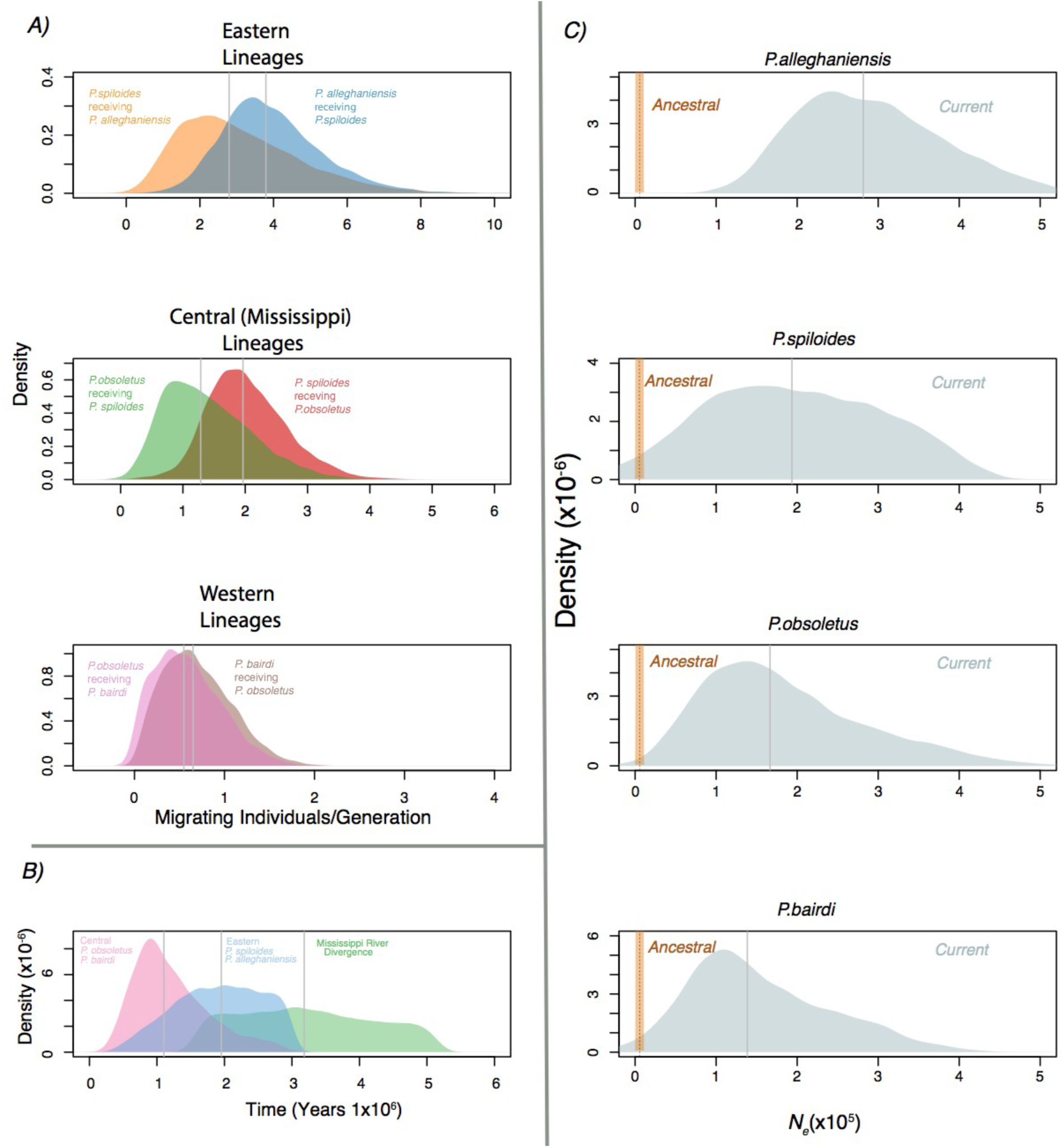
Results from PipeMaster showing estimates and directionality of migration (2Nm) (A), divergence times between taxon pairs and across the Mississippi River (*P. alleghaniensis/ P. spiloides* vs *P. obsoletus/P. bairdi*) (B), and changes in population sizes over time (C).

### Spatial population genetics and hybrid zones

We used ANN to determine what features of the landscape and environmental layers through time best predict genetic structure (Fig.4). Estimates of accuracy using ANN were high (>90%) for the first PCoA axis of genetic distance and declined rapidly thereafter approaching random shuffling of the response variable by axis 4-7 (Fig. 4). Most genetic structure was predicted by the Mississippi River showing 100% variable importance, when examining population structure across all four species. Spatial components alone also played a role in structuring these data for all species-pair comparisons, although they were partialed out in RDA analyses showing the significant effect of environmental and biogeographic variables (see Supporting Information Material 1). For those lineages east of the Mississippi River, population structure was predicted by climate at the Last interglacial and current environment for the first PCoA axis. For those lineages west of the Mississippi River, elevation, Last Glacial Maximum and eceoregions were important for structuring these lineages in the first PCoA axis. We note that while accuracy decreased on the 2^nd^ and 3^rd^ PCoA, other environmental variables still showed importance.

**Fig 4.**
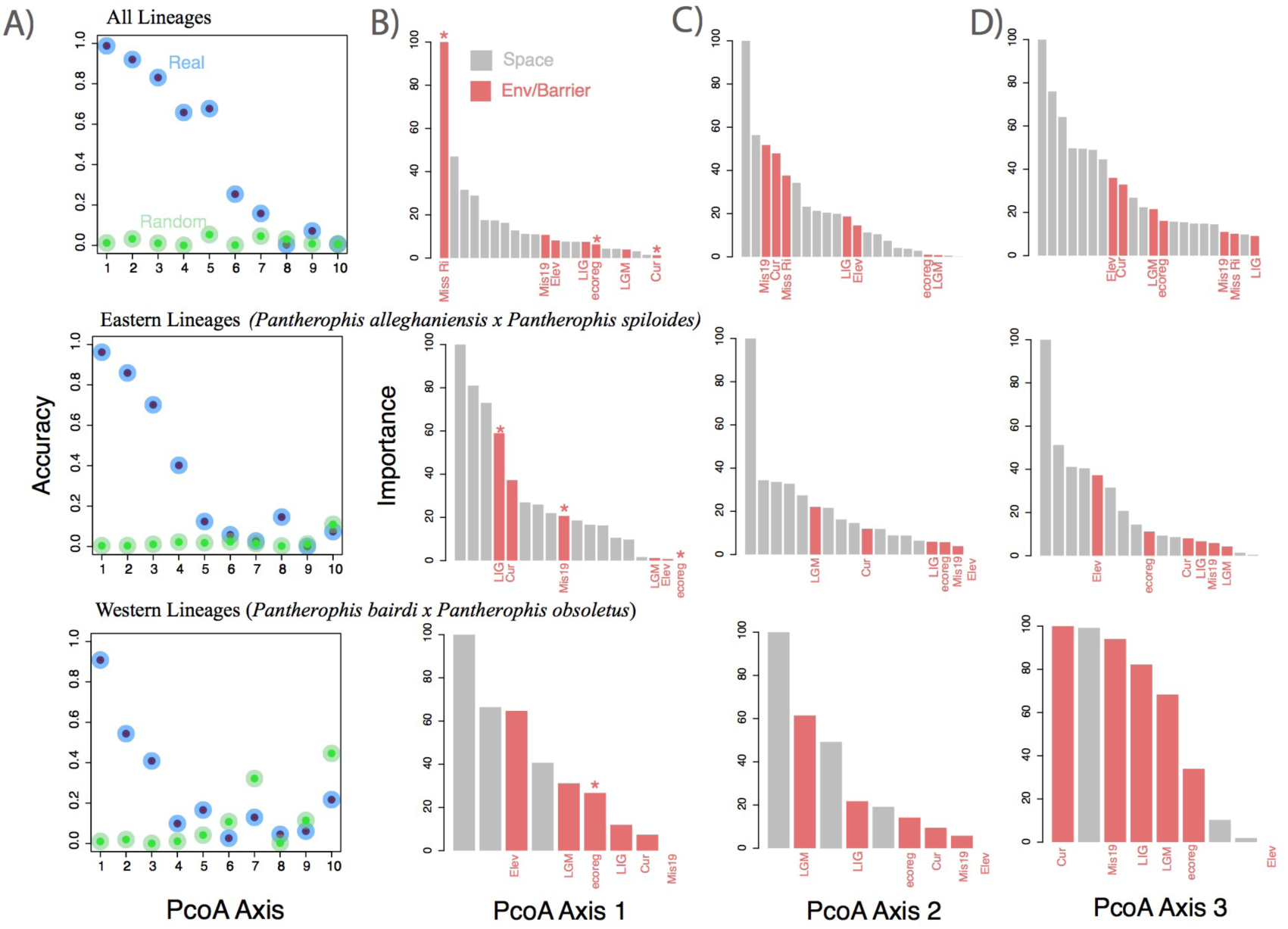
Artificial neural network results showing accuracy predicting genetic structure for 10 PCoA axes among all lineages and species pairs for real and randomly shuffled dependent variables (A). Variable importance are shown for spatial data (grey; principal coordinates neighbor matrices) and environmental charcteristics (red; B-D). Variable abbreviations are: Miss Ri (Mississippi River), Elev (elevation), ecoreg (ecoregions), Mis19 (Pleistocene Marine Isotope Stage 19 - 787 Kya), LIG (Last interglacial – 130 Kya), LGM (Last glacial maximum – 21Kya), and Cur (Current climate data). Asterisks above red columns are significant after corrected for space using redundancy analyses (RDA).

Using TESS admixture estimates and one-dimensional spatial distances, HZAR showed support for a model with no tails for all lineage-pair comparisons (eastern ΔAIC =-4 .08 − −8.34; central ΔAIC = −3.55 − −8.17; western ΔAIC = −2.12 − −8.48; Fig. 5, Table 1). Median width of clines decreased from the eastern lineages (256 km), to central (147 km) to the western lineages (21 km). Using environmental distances, both the eastern (ΔAIC = −4.39 − −9.45) and central lineages (ΔAIC = −4.32 − −346.38) showed support for a model with no tails for both, whereas the western lineages supported a left-tailed model (ΔAIC = −6.12 − −11.55). As a percentage of the entire environmental distance for a species-pair comparison, the environmental cline width decreased from the central (5.0%), eastern (4.0%), and western lineages (2.1%; Table 1). Percentages of widths per species-pair comparison were lower as a measure of total environmental distance than spatial distance.

**Fig 5.**
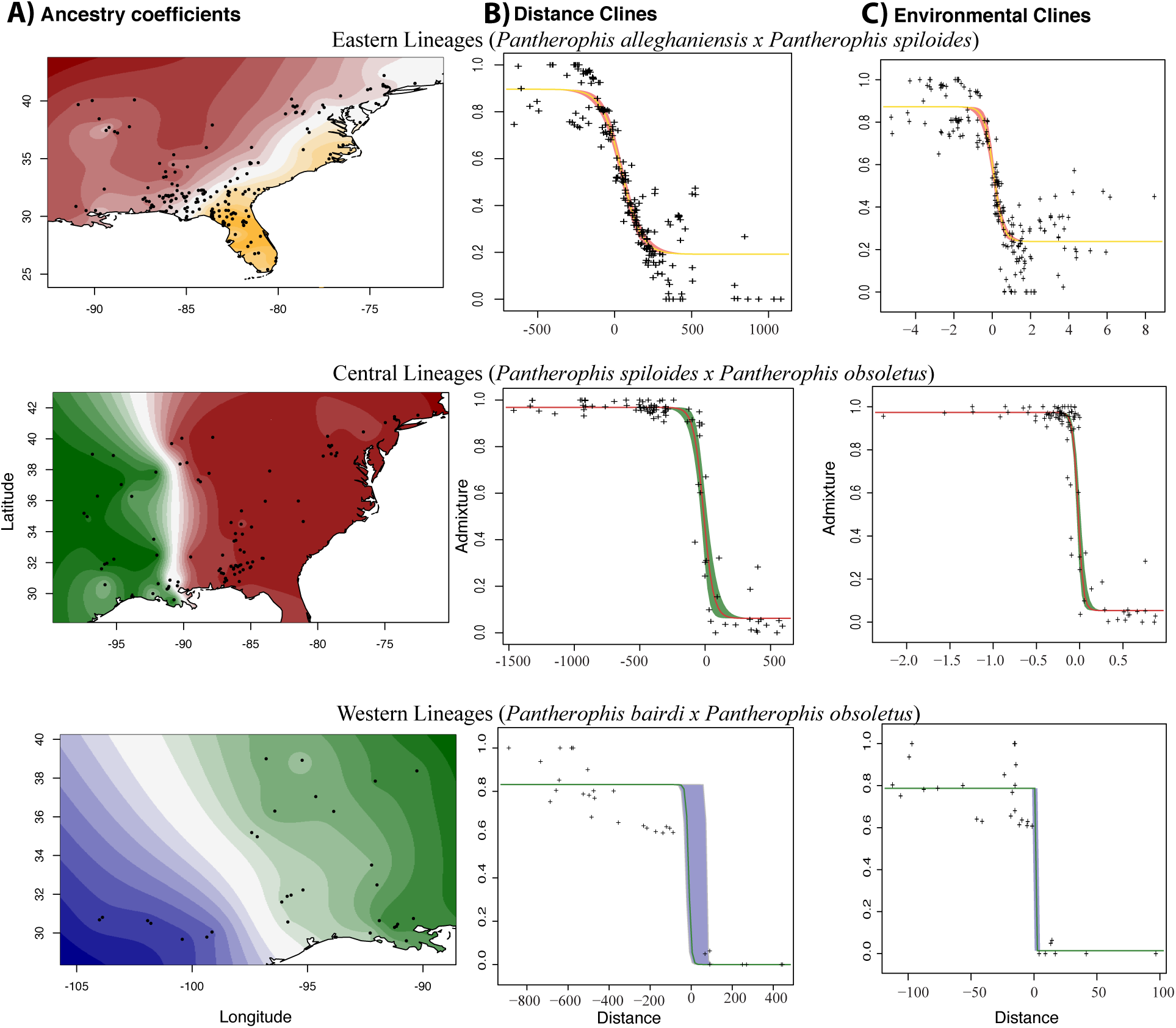
Using admixture from TESS3r (A), Gaussian cline (HZAR) model predictions are shown for both spatial (B) and environmental distances (C) for all three species pairs.

We examined cline widths by locus for each species pair (Fig.6). For each comparison a large number of loci were uninformative about cline widths, due to a lack of fixed differences among parental taxa. However, as a measure of expected cline width per locus under neutrality from the last glacial maximum, we showed that widths should be much larger (∼633-733 km) than found using admixture coefficients and across many loci (Fig. S5). We estimated that 128 loci between the central lineages, 8 between the eastern lineages, and 4 between the western lineages were both fixed in the tails of a cline for each parental taxon(ΔP>0.80) and presented cline widths lower than neutral expectations,(Fig. 6). Median cline centers for all loci, where the direction of selection on an allele was expected to change, was close to zero, the predicted center based on admixture proportions, for the central and western lineages, −8.32 and 4.95 km, respectively. However, center estimates ranged from −28.35 to 1766 for the eastern lineages, though a large number of loci (n=66) estimated the center closest to zero (ranging 20 km on each side; Fig. 6).

**Fig 6.**
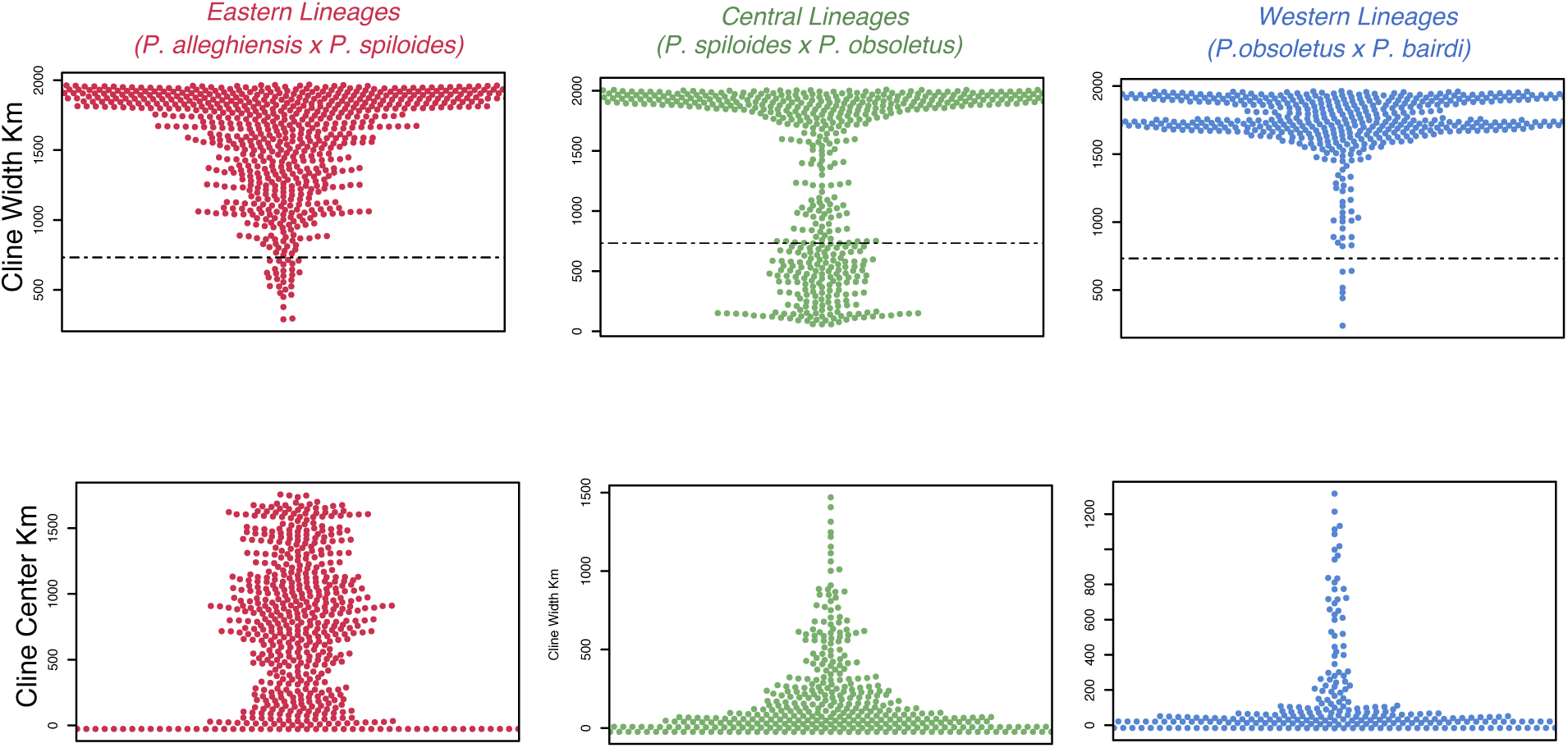
Beeswarm plots showing predicted cline widths and centers for all three species pairs over all loci using HZAR. Cline widths for loci lower than the predictions for neutrally forming hybrid zones since the last glacial maximum fall below the indicated dashed line.

When using a single SNP per locus we showed that 30% of loci from central lineages, 10.9% of loci for western lineages, and only 1.9% of loci for eastern lineages attained an F_st_ >0.35. Given demographic histories between comparisons, this threshold for F_st_ was generally considered extreme population disconnection beyond which advantageous loci will be exchanged between populations (Wright 1978; Frankham et al. 2010).

## Discussion

### Grey Zone Dynamics

To understand the factors shaping reproductive isolation in a species complex, we show that divergence time does not predict rates of migration or cline width across space in eastern ratsnakes. While reproductive isolation should increase with time (Orr 1995; Singhal and Moritz 2013), our results indicate that overall genomic estimates of gene flow may be inadequate to assess reproductive isolation with time. In contrast, the number of loci showing steep cline shifts in contact zones is related to timing of divergence. Disconnection between episodes of repeated secondary contact through time differing among species-pair comparisons or variable strength of ecological divergence may account for the dissociation between time and overall migration rate.

Using multiple methods to examine genetic structure and migration over space and accounting for isolation by distance, we show that the eastern ratsnakes are composed of four divergent lineages, with *P. alleghaniensis* and *P. spiloides* meeting in the southeastern US at the intersection of subtropical and temperate areas, *P. spiloides* and *P. obsoletus* separated at the Mississippi River, and *P. obsoletus* and *P. bairdi* meeting at the connection on the forested and rocky habitats on the western edge of the Edwards Plateau. The western lineages (*P. obsoletus*, *bairdi*) have low migration rates, a cline equivalent to only 1.6% of the parental range, large fixation indexes (F_st_ > 0.35) in 10.9% of loci, and likely geographic selection in only 4 loci. This is in comparison with central lineages (*P. obsoletus, P. spiloides*) showing more migration, a cline occupying 7.2% of the parental ranges, but with 35% of loci showing high F_st_, and geographic selection in 128 loci. Finally, eastern lineages (*P. alleghanienis, P. spiloides*) have the highest rate of migration, with only 1.9% of loci showing high fixation, and only 8 loci revealing geographic selection (Figs. 3 & 6). The oldest species-pair at the Mississippi River have moderate migration/generation but yet have the highest number of loci showing steep clines at the hybrid zones. Similar to other studies where many loci showing differential introgression or variance in cline slope (Nosil et al. 2009; Carneiro et al. 2013; Stankowski et al. 2019), our results lend support that the degree of reproductive isolation may be defined at the level of the genes and not the genome (Macholán et al. 2011; Harrison 2012). Therefore, it is likely that these species-pairs of ratsnakes are traversing different phases of reproductive isolation, from the genic to genomic, and therefore overall coalescent estimates of migration alone may not useful proxies for understanding position in the gray zone.

Hybrid zone dynamics help to understand the interactions between pairs of taxa as characteristics of the hybrid zone reveal whether there is selection against hybrids (tension zones), selection for hybrids (bounded superiority), or ecological gradients with reduced fitness. They also reveal whether selection is endogenous (genomic) and/or exogenous (environmental; Barton and Hewitt 1985; Barton and Gale 1993). Our results that combine estimates of when and how divergence occurred with contemporary dynamics of hybrid zones provide a comprehensive view of how lineage structure is maintained through time. Demographic models for ratsnakes indicate that some migration at the Mississippi River and among all species pairs occurred during the Pleistocene suggesting primary or secondary contact existed through major climate change cycles. Population expansion in all lineages occurred at a minimum before the LGM, likely indicating ancient contact and formation of these hybrid zones. Given this age and assuming selective neutrality, we demonstrate that even after the last glacial maximum (LGM) that these hybrid zones should be much larger, in most cases exceeding the ranges of parental taxa (Fig. S5). The widths of the current hybrid zones have likely been maintained or continuously reformed through time after reemerging from glacial refugia, as discussed for other temperate taxa (Hewitt 1996, 2011). Despite constant connectivity, the historical and geographic identity of the parental taxa are preserved. While there is likely some selection against hybrids as estimated from hybrid zone width and dispersal, depending on species-pair comparison, it is possible that other factors may be constraining these hybrid zones, for example fitness changes along an ecological gradient or isolation by environment (Case and Taper 2000; Case et al. 2005; McEntee et al. 2018). For all groups, our ANN show that either past or current environmental differences or barriers to dispersal are important for structuring overall genetic differences among species pairs. Among contemporary lineages and analyses, we show that admixture clines change sharply at environmental transitions relative to just spatial distances (Fig. 5), which suggests that fitness is likely changing across these ecological areas. Similarly, we find several loci in each species-pair comparison that show similar or smaller cline shifts in these areas found for overall admixture predictions. Unless these loci represent the outcomes of isolation by distance and fixation in each ends of the cline, there is likely some selection on these genes. Therefore, barriers that isolated these lineages initially, along with combinations of current and past environmental changes across the ranges of all these species pairs, likely helped form and maintain these hybrid zones.

### Influence of barriers and environment on population structure

These ratsnake lineages are structured by a complex of physical barriers, historical climate profiles, and geographic distance. Machine learning approaches show that the Mississippi River has served as a strong barrier for isolating lineages of ratsnakes, forming groups east and west of this river during the late Pleistocene/Pliocene (Figs 1&3,4, Table 1). These results are consistent with previous mtDNA and morphological conclusions (Burbrink, 2001; Burbrink et al., 2000). The Mississippi River has served as a strong barrier to gene flow given evidence from historical estimates of increased drainage (Cox et al. 2014) and numerous unrelated organisms show genetic disconnection across the river (Robison 1986; Burbrink 2002; Soltis et al. 2006; Burbrink et al. 2008; Pyron and Burbrink 2009; Brandley et al. 2010; Satler and Carstens 2017; Myers et al. 2020). Hybridization, while low (1-2 individuals/generation, Fig.3), does occur in a zone (∼100km) east of the Mississippi River between the central lineages (*P. obsoletus* and *P. spiloides*). Furthermore, genetic structure is also predicted by geographic distance and mid-Pleistocene climate change (787kya).

The Mississippi River is responsible for the earliest divergence in this complex but other geographic features are also important. East of the Mississippi River, divergence between *P. alleghaniensis* and *P. spiloides* was influenced by climate at the last interglacial, current climate, and geographic distance (Fig 4). For these taxa, divergence occurred at the Appalachian Mountains and Apalachicola/Chattahoochee River System. *Pantherophis alleghaniensis* may have diverged in an isolated Florida, due to sea level changes during interglacials, as suggested by our ANN, and subsequently expanded north and west. This isolation is reflected by the unique morphology there showing a high concentration of the yellow, striped color patterns in the range of *P. alleghaniensis* in Florida and parts of the southeastern coast of the US (Schultz 1996; Burbrink 2001). This suture zone between the Florida peninsula and the continental US has been found for many taxa (Remington 1968; Swenson and Howard 2005; Burbrink et al. 2008). While these two forms can be delimited with genetic data and occupy distinctly different geographic regions, reproductive isolation may not be complete. Although, individual loci likely show varying levels of reproductive isolation. Niches are also different between these taxa; *P. alleghaniensis* is typically found in the Florida peninsula and coastal plains environments (Bailey 1995; Burbrink 2001), whereas *P. spiloides* is found throughout the remainder of the forested habitats east of the Mississippi River including the ecoregions defined as interior river valleys and hills, interior plateau, Appalachian habitats, southeastern and southcentral plains, and Mississippi River valley. Uncertainty in the identity of taxa using morphology where these taxa meet is likely due to extensive gene flow in hybrid zones at barriers following postglacial range expansion. Finer-scale testing is necessary to tease apart the effects of these river systems, ancient embayments, and connections to uplifted areas. Although the timing of divergence predates the last interglacial, it is possible that repeated 100,000 year glacial cycles isolated these taxa into refugia, as has been suggested previously for ratsnakes and other organisms (Fig.3; Waltari et al. 2007; Noss et al. 2015).

Divergence between *P. bairdi* and *P. obsoletus* occurred in the western edge of the Edwards Plateau, with the former extending westward into xeric forested or Chihuahuan desert scrub associated with rocky habitats, whereas the latter extends east into Texas, preferring forested habitats, (Lawson and Lieb 1990; Werler and Dixon 2000). Population genetic structure between these taxa is associated with elevation, the last glacial maximum, ecoregion differences, and geographic distance (Fig. 4). Dates of origin of these taxa are within the mid-Pleistocene, suggesting that past climate played a strong role in promoting population divergence. During this period, the Chihuahuan Desert was much drier relative to the late-Pleistocene (Graham and Mead 1987; Metcalfe et al. 2002; Metcalfe 2006) and this increased aridity could have driven ecological divergence between this species pair. While divergence time is not the oldest here, migration rates are lowest, and the hybrid zone is narrow. Previous research shows limited hybridization with backcrosses between these taxa occurring in a narrow region of the southwestern Edwards Plateau (Lawson and Lieb 1990; Vandewege et al. 2012).

Despite each lineage occupying uniquely different habitats, population responses to Pleistocene climate change were similar (Fig. 3). Given the divergence time for each group, *N_e_* has expanded synchronously by 50 times since the Pleistocene. These estimates are consistent with other predictions in the ENA, where 75% of the tested vertebrates show population size expansion and 75% of tested snakes show synchronous expansion of *N_e_*, likely coinciding with the retreat of glaciers at various times in the Pleistocene or during the Holocene (Bintanja and van de Wal 2008; Burbrink et al. 2016). However, *Pantherophis bairdi* has a range only in a small area of the west Texas and isolated populations in northeastern Mexico, where habitats may have been indirectly affected by glacial cycles.

### Taxonomy and delimitation in the genomic age

Delimiting species using genetic data, while useful, has proved controversial (Bauer et al. 2011; Sukumaran and Knowles 2017; Leaché et al. 2019). In particular, most coalescent delimitation methods fail to account for gene flow (but see Flouris et al. 2019), do not consider spatial information, and place individuals into predefined groups prior to testing (O’Meara 2010). These methods all assume species diverge at an evolutionary time scale where mutations accumulate through drift or selection (Rannala 2015). And while simulations show that a threshold above one migrating individual/10 generations will cause model-based delimitation methods to fail at finding two species (Zhang et al. 2011), it is not clear how pulses or inconsistent migration affect species transitioning through the gray zone. Importantly, Wright (1931) indicated that up to 1,000 individuals migrating/generation may not prevent populations from drifting given effective population size and spatial segregation. Also, this does not consider the effects of selection and recombination producing variable introgression among loci.

Modern thresholds for species delimitation (Jackson et al. 2017; Leaché et al. 2019) use GDI (>0.7), estimates of species divergence time over population size (2τ/θ>1), absolute divergence time (10^4^) and maximum migration rates (M=Nm<1), metrics that are all likely influenced by sampling effort, location of samples sequenced, and detection of hybrids. For example, if large hybrid zones exist but remain poorly sampled then estimates of migration will be reduced, GDI will be high, and the probability of MSC estimators will suggest multiple species. Conversely, if only hybrid zones are sampled then all metrics will predict only a single species. For all pairs of ratsnakes 2τ/θ>1 and divergence time is greater than 10^4^. However, overall migration rates remain high between adjacent pairs in the eastern lineages. This forces us to address how we are defining a species in the context of hybrid zones, how we infer reproductive isolation beyond MSC methods, and how selection is maintaining hybrid zones. For all lineages of ratsnakes, hybrid zones are smaller than predicted by neutral selection if these hybrid zones formed in the Pleistocene. However, geographic lineage identity has been maintained for >133,000 generations in all pairs, regardless of migration. Moreover, where hybrid zones extend over large areas, gene flow likely varies throughout space and may have changed over time (Barton and Hewitt 1985; Mallet 2007), particularly in response to glacial cycles. Thus, identifying and delimiting species in the context of genetic admixture alone may be difficult without understanding the history of gene flow, and as shown here, specific locus introgression.

We have demonstrated that these four species, delimited previously using mtDNA and morphology (Burbrink, 2001; Burbrink et al., 2000) can be recognized as distinct taxa given methods of spatial grouping and species delimitation, though ranges in the eastern-most lineage are restricted further south than previously thought. However, migration rates between *P. spiloides* and *P. alleghaniensis* (Fig. 3) may be high enough that they do not represent independent evolutionary trajectories (de Queiroz, 2007; Rannala, 2015; Zhang et al., 2011), though differential introgression among some might suggest that reproductive isolation is maintained for some loci. Unique niches in the subtropical areas of the southeastern US and morphology provides some evidence for isolation between these taxa: the striped forms (with variable ground colors of orange, yellow, green, and gray) are indicative of *P. alleghaniensis* whereas adults with dark saddle patterns on grey, olive, brown, or black ground colors (or completely black with no pattern) are found west of Florida and east of the Mississippi River, throughout the Midwest and up to the northeastern US and Canada represent *P. spiloides*. However, hybridization north and west of Florida is likely represented by a dulling of this yellow ground color and mixed patterned individuals (Schultz 1996; see Supporting Information Material 1 for more information on taxonomy).

Modern genomic tools have advanced our understanding of timing of divergence, population isolation, lineage connectivity, and the nature of hybrid zones. As we show with the eastern ratsnakes, we can better comprehend processes of speciation by integrating studies of hybrid zones to understand changes in admixture and differential introgression with coalescent estimates of isolation and migration. We also show that timing, rates of gene flow, and differential introgression at hybrid zones are unique to each species pair, likely traversing different genic and genomic scales of divergence in the gray zone across changing environments in the Eastern Nearctic. Future studies should examine clines with more detailed sampling using whole genomes to connect weakly introgressing loci with traits.

## Acknowledgments

We thank R. Glor, R. Brown, and L. Welton at KU and C. Austin and D. Dittman at LSUMN and K. Krysko and FLMNH, D. Shepard at Louisiana Tech University, and A. McKelvy for providing tissues. We also thank L. and G. Derryberry for help with HZAR. We also thank E. Chen for help extracting DNA. EAM was funded by the Gerstner Scholar/Theodore Roosevelt and Peter Buck/Walter Rathbone Bacon fellowships. We thank D. Frost, A. Leaché, A. Bauer, and K. deQuieroz for helpful discussions on species delimitation, type specimens, and admixed type specimens.

## Data Accessibility Statement

All data and unique scripts are available on Dryad XXX.

## Supplemental Material

### Supporting Information Material 1

### Supplemental Methods and Results

#### Dataset

We sampled 288 individuals liberally covering the range of all species within the *Pantherophis obsoletus* complex (Fig.1; Dryad XXX). DNA was extracted from all samples using Qiagen DNeasy Blood & Tissue Kits and samples were screened for quality using broad-range Qubit Assays (https://www.thermofisher.com/us/en/home/industrial/spectroscopy-elemental-isotope-analysis/molecular-spectroscopy/fluorometers/qubit/qubit-assays.html). We used services from RAPiD Genomics (https://www.rapid-genomics.com/services/) to generate 5472 baits and to sequence 5060 conserved elements (UCEs) loci following the protocols from (Faircloth et al. 2012) and (Sun et al. 2014).

Raw sequence reads were trimmed of adapter contamination using illumiprocessor (Faircloth 2013), a wrapper around the trimmomatic package (Bolger et al. 2014). These sequence capture data were then assembled by mapping reads to all UCEs found within a Chromium 10x assembled *Pantherophis spiloides* genome (Burbrink and Myers, *in prep*) after 2,500 base pairs of flanking DNA around UCE baits were pulled from this genome assembly using the Phyluce commands *phyluce_probe_run_multiple_lastzs_sqlite* and *phyluce_probe_slice_sequence_from_genomes* (Faircloth 2016). Sequence reads for each sample were then mapped to the ‘reference’ UCE dataset using bwa v0.7.17 (Li and Durbin 2009).

Samtools was used to convert sam format to bam files and alignments were soft clipped using the CleanSam tool in Picard (http://broadinstitute.github.io/picard/). To phase these data, we mapped reads back to each individual sample’s alignments. To do this, bam files were converted to fastq using samtools mpileup (Li et al. 2009), fastq files were converted to fasta format via seqtk (https://github.com/lh3/seqtk) and all missing sites were removed using seqkit (Shen et al 2016). These newly created fasta files were then used as a reference sequence for phasing the data following the seqcap_pop pipeline (Harvey et al. 2016). Briefly we followed this pipeline and used GATK (McKenna et al. 2010) to mark duplicates, call, realign, and annotate/mask indels, call and annotate SNPs via GATK, restrict SNP calling to high quality SNPs, and then conducted read backed phasing. Finally, we created a sample specific fasta file of all phased loci for every sample using ‘add_phased_snps_to_seqs_filter.py’ available via the seqcap_pop pipeline (Harvey et al. 2016). All samples were combined into locus specific fasta files using a perl script developed in (Myers et al. 2019). Locus specific fasta files were aligned using muscle as implemented in the R package ape (Paradis and Schliep 2019) after removing individuals from alignments that fell below the 25% quantile of the distribution of sequence lengths for each locus. All fasta files were conservatively trimmed of missing data resulting in alignments with no missing sites (i.e., trimming to the shortest sequence in the alignment). We filtered these alignments by removing all loci that were missing >50% sequenced samples and removed all individuals that were missing from >30% of all alignments. Finally, we created a vcf file from these alignments and using vcftools (Danecek et al. 2011) and filtered this for minor allele frequency >10% retaining only 1 SNP/locus for population assignment analyses (Linck and Battey 2019).

#### Geographic groupings

We estimated population structure by comparing Discriminant Analysis of Principal Components (DAPC; Jombart et al., 2010) and spatial Principal Component analysis (sPCA; Jombart et al., 2008) in adegenet v2.1.2(Jombart 2008), sparse nonnegative matrix factorization (SNMF; Frichot et al., 2014) in LEA v1.4.0 (Frichot and François 2015), and estimated effective migration surfaces (EEMS v; Petkova et al., 2016), which accounts for IBD. Each of these methods use different assumptions for grouping individuals and we compare congruence among them. The model-free method DAPC sequentially estimates K-means for clustering and selects the best fit to infer genetic clusters in the absence of prior group identification. Using the package adegenet (Jombart 2008) in R (R Core Team 2015) we first transformed data using PCA by choosing a large number of PC axes (n=200), then picking the number of discriminant functions yielding large F-statistics (>2000) and optimal number of groups using BIC (Bayesian inference criterion). Because using a large number of the PCs may yield arbitrary solutions, we estimated the optimal *a-score*, which measures this bias by calculating the difference between the actual cluster assignment probabilities and randomly assigned probabilities (predicting 15-23 PCs). We also used cross-validation with a 90% training and 10% test dataset to estimate the average predicted success for each group.

Because DAPC does not account for spatial structure, such as IBD, we also used sPCA. Similar to DAPC, this method does not require data to be in Hardy-Weinberg or linkage equilibrium. sPCA also adds an element of space by isolating global structure, representing disconnected groups or clines, from local structure, representing repulsion (selection against similar genetic types co-occurring), and random noise. Using adegenet, we estimated sPCA given SNPs and locations of each sample using up to 15 positive eigenvalues (global structures) and 15 negative eigenvalues (local structures). To connect individual locations to build a connection network, we used Delaunay triangulation. We also implemented Monte-Carlo tests with 100 permutations using the function *spca_randtests* to determine if significant local or global spatial structures exist.

We also assessed ancestral coefficients using SNMF in the package LEA (Frichot and François 2015). This methodology is comparable to ADMIXTURE (Alexander et al. 2009) and Structure (Pritchard et al. 2000) methods, but runs 10-30x faster using genomic-scale data. This method produces a least-squares estimate of ancestry populations given *K* ancestral populations We estimated ancestry coefficients over 1-6 *K* using 100 repetitions per *K*, 100 iterations per algorithm, and a tolerance of 0.05. To determine the appropriate number of groups, we used a cross-validation technique with the entropy criterion, which masks genotypes to fit a particular model to each *K*. Finally, to compare these methods of grouping individuals into populations, we estimated migration rates over the landscape using EEMS (Petkova et al. 2016). This method models effective migration rates over geography to represent regions where migration is low in cases when genetic dissimilarity increases rapidly, thus providing a uniquely distinct view of the location of population clusters relative to biogeographic barriers as compared to DAPC, sPCA, and SNMF. We ran EEMS 3 times for 3x10^6^ generations with burnin set at 1x10^6^ generations, thinned by 9,999 generations, with 1,000 demes covering the known range of the *P. obsoletus* complex (Burbrink 2001).

Both EEMS and DAPC find population structure, generally matched the geographic ranges of *P. alleghaniensis*, *P. spiloides, P. obsoletus*, and *P. bairdi* (Fig. 1 and Fig. S1; Burbrink, 2001). Total DAPC assignment probabilities were 0.975 (*P. bairdi* = 1.0, *P. obsoletus* = 0.96; *P. spiloides* = 1.0, and *P. alleghaniensis* = 0.97). Comparing these assignments to those from SMNF, we found that only *P. alleghaniensis* was misclassified as *P. spiloides* 8.5% of the time. Similarly, SNMF cross validation predicted four groups (cv = 0.599) as compared to two (cv=0.627), three (cv=0.607), and five (cv=0.601). These four groups generally match the same ranges as those found in DAPC. sPCA showed the presence of four groups, with PC1 showing the separation of groups at the MR and the two lineages east of this river, whereas PC2-5 shows divergence in all taxa. Importantly, global spatial structures were significant (observed spatial structure test statistic = 75.47, random expectation = 28.14; *P* < 0.009). Similarly, all three EEMS runs suggest low migration at the MR and the area separating the *P. bairdi* and *P. obsoletus* in west Texas. East of the MR, estimated low migration occurred near the Appalachian Mountains, though this area is a complex mixture of isolation and gene flow (Fig. 1). All three EEMS generated similar acceptance proportions for all proposal types (12-51%).

#### Running SNAPP

We set forward and reverse mutation rates to 1. We applied a gamma distribution (α = 2, β =200) for the speciation rate (λ) prior and the *snapprior* was set to α = 1, β =250, and κ = 1. To test the four alternative species delimitation models including four taxa, three taxa (collapsing *P. bairdi/P. obsoletus* or *P. alleghaniensis/P. spiloides*), and two taxa (collapsing *P. bairdi/P. obsoletus* and *P.alleghaniensis/P. spiloides* we used a stepping-stone analysis with the *PathSampleAnalyser* with 72 steps and a chain length of 200000, burnin percentage at 50%, and a preBurnin of 10,000. We checked for stationarity using Tracer v1.7.1 (Drummond and Rambaut 2007) determining that estimated sample size (ESS) were > 200 for all parameters.

#### Running PHRAPL

PHRAPL uses gene-tree distances between simulated and empirical data to rank models using approximate likelihoods and AICs. To generate the empirical observations we estimated unrooted gene trees for each loci using IQ-Tree 1.6.12 (Nguyen et al. 2015) and testing all substitution models per locus prior to estimation. We rooted the trees using midpoint rooting and performed 100 subsamples of two individuals per population. To reduce model space, we allowed only one population size parameter (all populations have equal size) and one migration parameter (all migration rates are equal). We then ran 1000 simulations per model, searching over a grid of four CollapseStarts values and five MigrationStarts values.

#### Spatial genetics using RDA

We predicted genetic distances from environmental, elevation, river, ecoregion, and distance variables using RDA in the R package Vegan (Dixon 2003), which avoided problems with spatial data using Mantel tests (Legendre et al. 2015). This method was used to compare results from the ANN analyses. RDA is an asymmetric canonical analysis here used to determine the significance of predictor axes (Legendre et al. 2011) at generating predict genetic structure over a landscape (McGaughran et al. 2014; Noguerales et al. 2016). We converted spatial distances using principal coordinates of neighbor matrices (PCNM) in the r package vegan (Dixon 2003) keeping all axes with positive eigenvectors. We ran this using the *capscale* function with only spatial variables alone to determine which significantly predict genetic distance to be passed to out ANN analyses. We also ran this with the significant spatial axes partialed out and environmental, elevational, ecoregion, and river variables. All variable distances were used to predict genetic distance and significance of each variable was tested using anova-like permutation tests to examine the joint effect of predictor variables given partial effects from spatial distance to account for IBD.

Estimates of accuracy using ANN were high (>90%) for all comparisons and yielded similar conclusions to RDA in that environmental and geographic barriers and not just spatial variables predicted genetic structure. Here, RDA demonstrated that most of the structure for all taxa can be predicted by the Mississippi River (*P*=0.001), ecoregion (*P* =0.008), and current climate (*P*=0.019). Distance also played a role in structuring these data, though they were partialed out in RDA. For those lineages east of the Mississippi River, population structure was predicted by ecoregion (*P*=0.014), Pleistocene Marine Isotope Stage 19 (787 Kya; *P*=0.001), last interglacial (*P*=0.007) and ecoregion (*P*=0014). For those lineages west of the Mississippi River, we demonstrated predicted by ecoregion (*P=* 0.001).

#### Genomics, the ICZN, and Taxonomy

One problem likely to be encountered by other researchers using genomic data is that previously named type specimens at particular type localities may be from regions that show some admixture. For instance, the type locality of the eastern-most species, *P. alleghaniensis* is “the summit of the Blue Ridge in Virginia and the Highlands of the Hudson” (Holbrook 1836; Schultz 1996). Both areas are likely composed of admixed individuals and the International Code of Zoological Nomenclature (ICZN) forbids naming species based on hybrids (Article 1.3.3;ICZN, 1999) though the intention of this ICZN article did not consider proportions of admixture. One possible solution would be to use the next available name from Florida, where no admixture is present (eliminating *P. quadrivittata*), which would be *P.deckerti* (type locality: Lower Matecumber Key, FL; Brady 1932). Another solution would be to designate a Neotype from a locality showing no admixture and retain *P. alleghaniensis.* If *P. alleghaniensis/P. spiloides* were not considered unique, then *P. alleghaniensis* has priority by 18 years (Holbrook 1836; Duméril et al. 1854).

The type locality for the remaining taxa, *P. spiloides (*New Orleans, Louisiana; Duméril et al. 1854), *P. obsoletus* (On the Missouri River from the Vicinity Isle au Vache to Council Bluff; Say, 1823), and *P. bairdi* (Fort Davis, Apache Mountains, Jeff Davis Co.: Texas; Yarrow 1880) are all from regions that likely represent individuals not admixed (Fig.1 & 5, Burbrink, 2001). Species delimitation methods, estimates of fixation for loci, and observation that these four organisms have remained geographically distinct in the face of gene flow throughout the Pleistocene suggests that these taxa should be considered unique species though hybrid zones exist. We acknowledge difficulties recognizing the eastern lineages as distinct and could argue for recognizing them as a single taxon, *P. alleghaniensis*.

---

**Fig S1.**
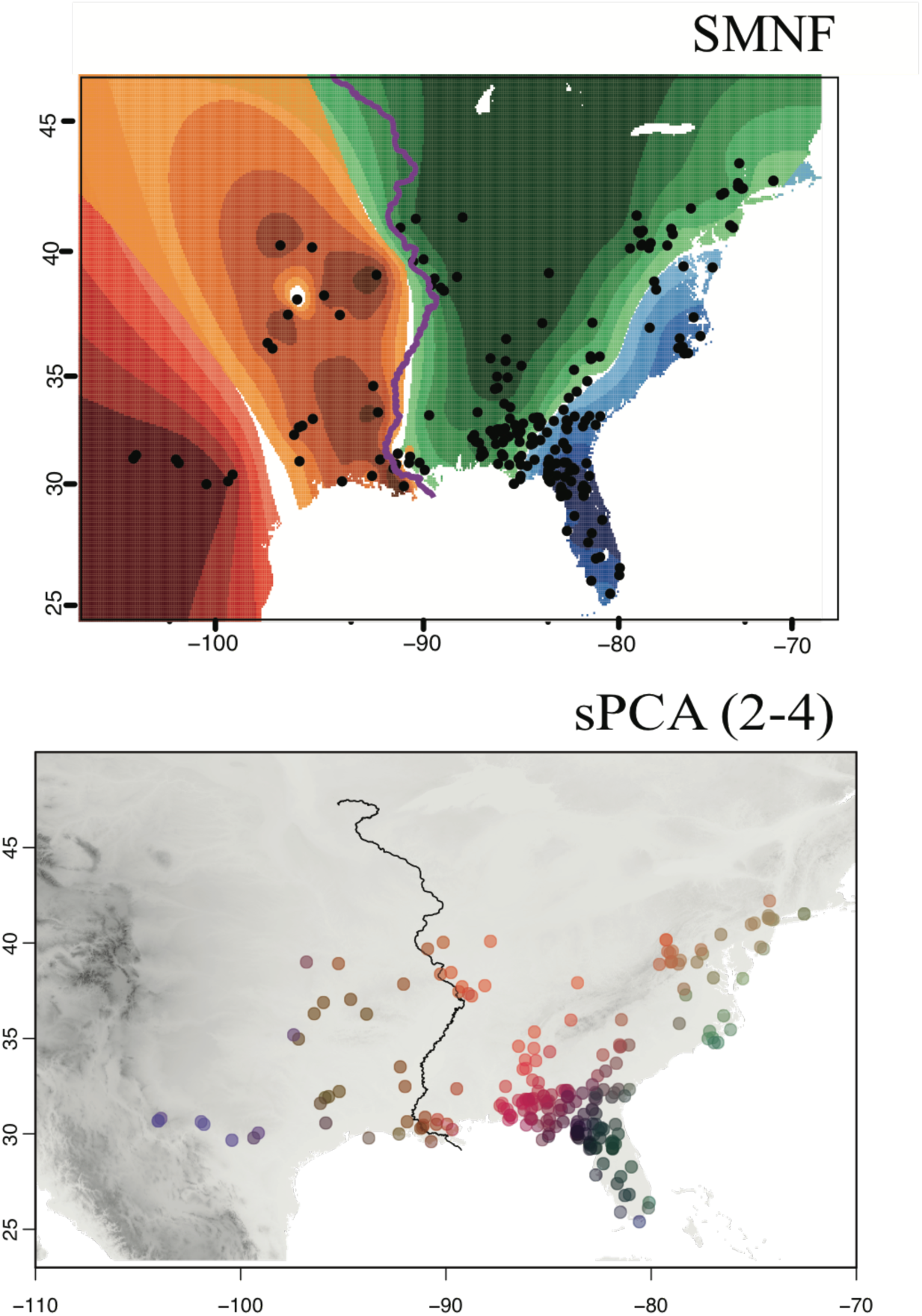
Delimiting populations using sparse nonnegative matrix factorization (SNMF) and spatial Principal Component analysis (sPCA) showing the location of four geographically distinct lineages.

**Fig S2.**
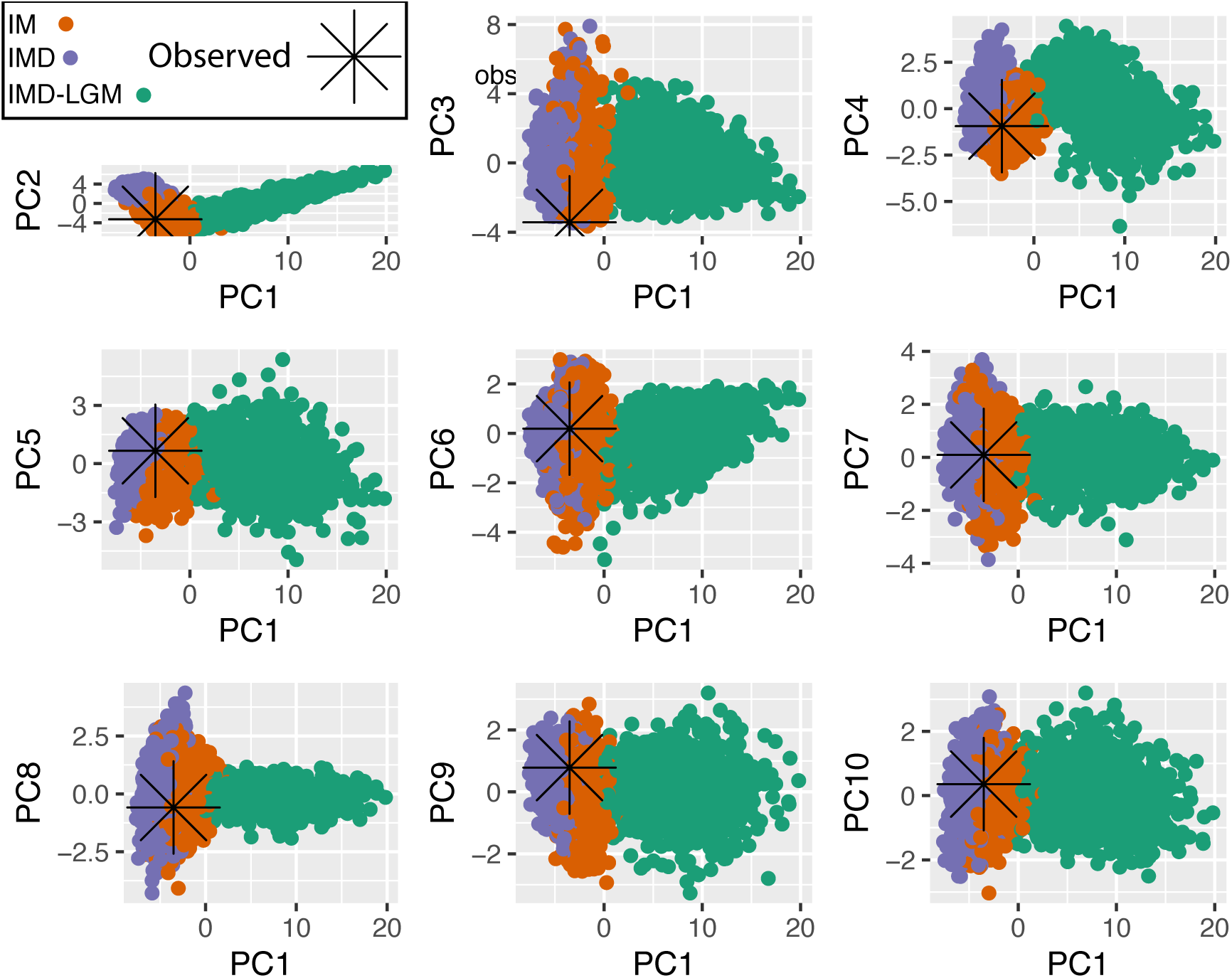
PipeMaster model fit of observed data to simulations under IM, IMD, and IMD-LGM models for PC1-PC10.

**Fig S3.**
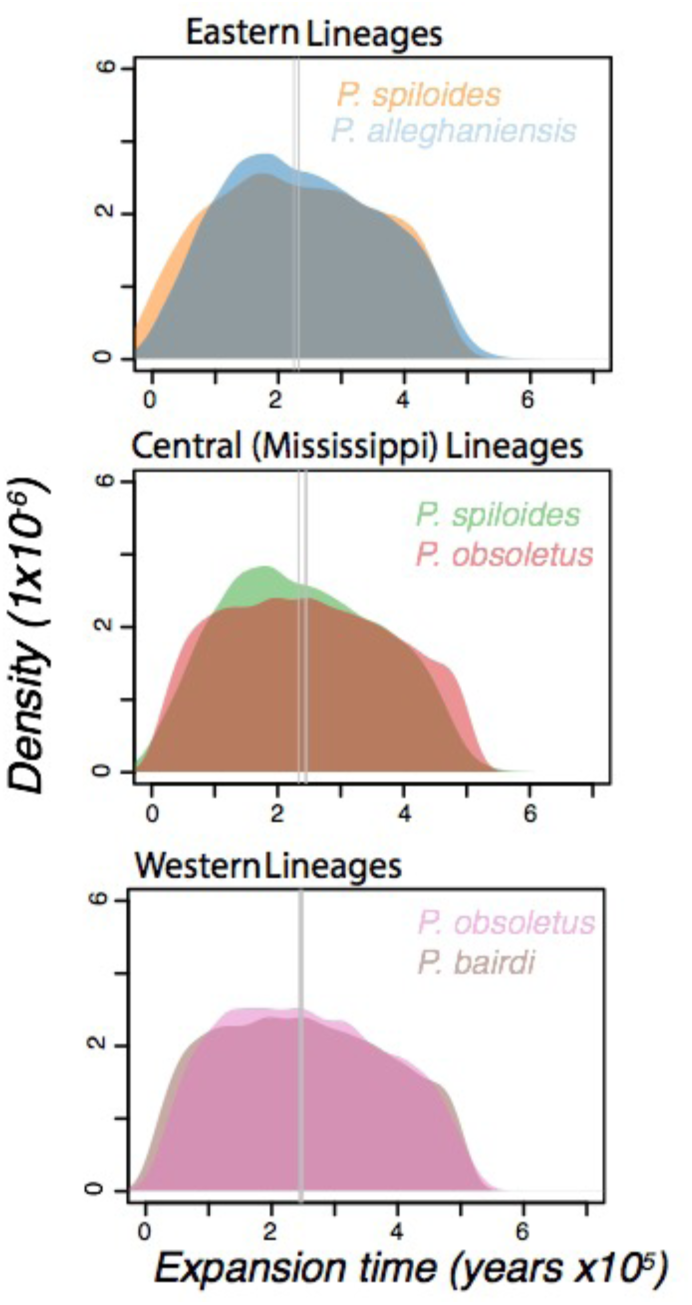
Expansion times estimated from PipeMaster for all adjacent taxon pairs.

**Fig S4.**
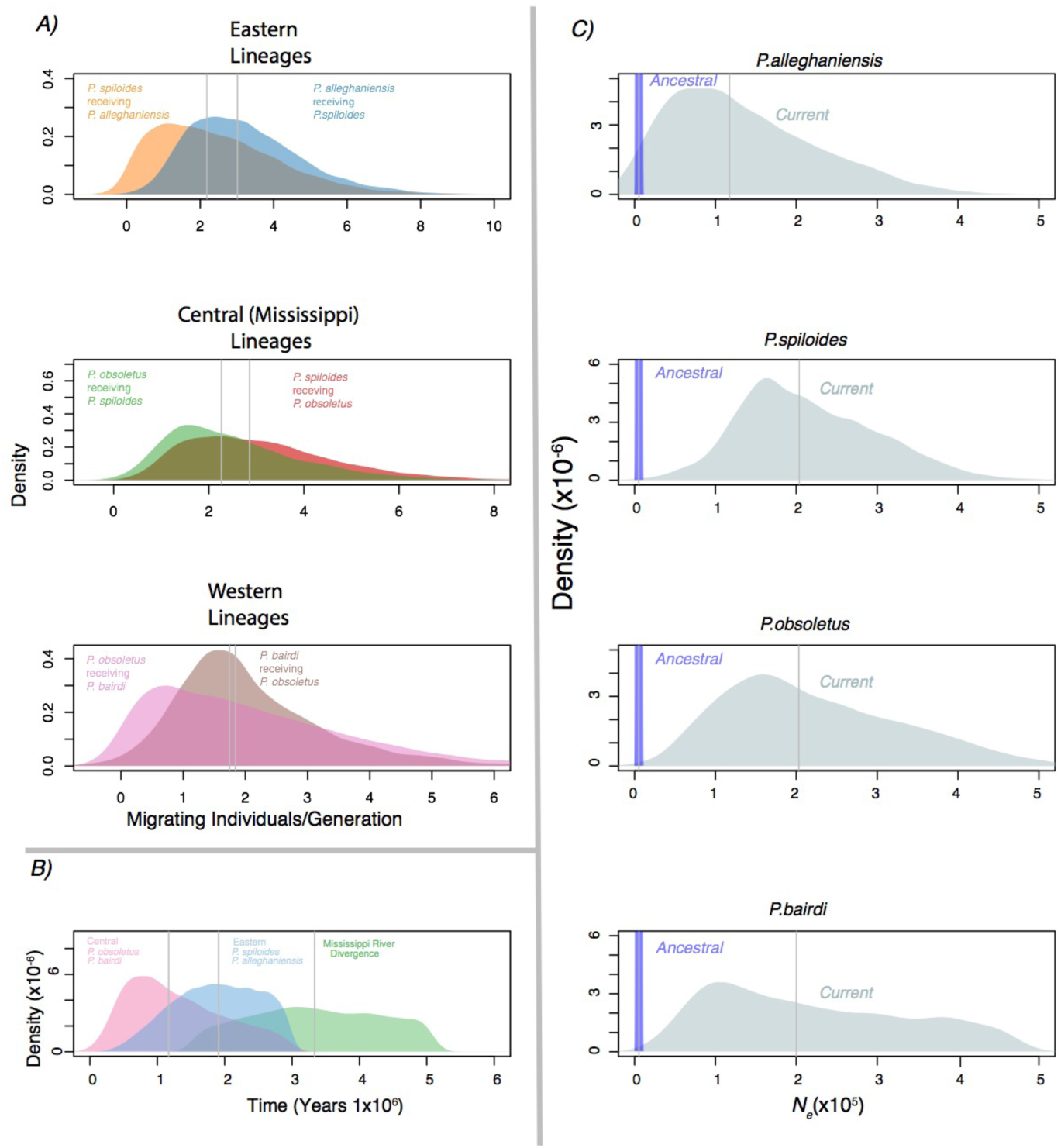
Results from PipeMaster after removing hyrbids showing estimates and directionality of gene flow (A), divergence times between groups and taxon pairs and the Mississippi River (*P.alleghaniensis/ P.spiloides* vs *P.obsoletus/P.bairdi*) (B), and changes in population sizes over time (C).

**Fig S5.**
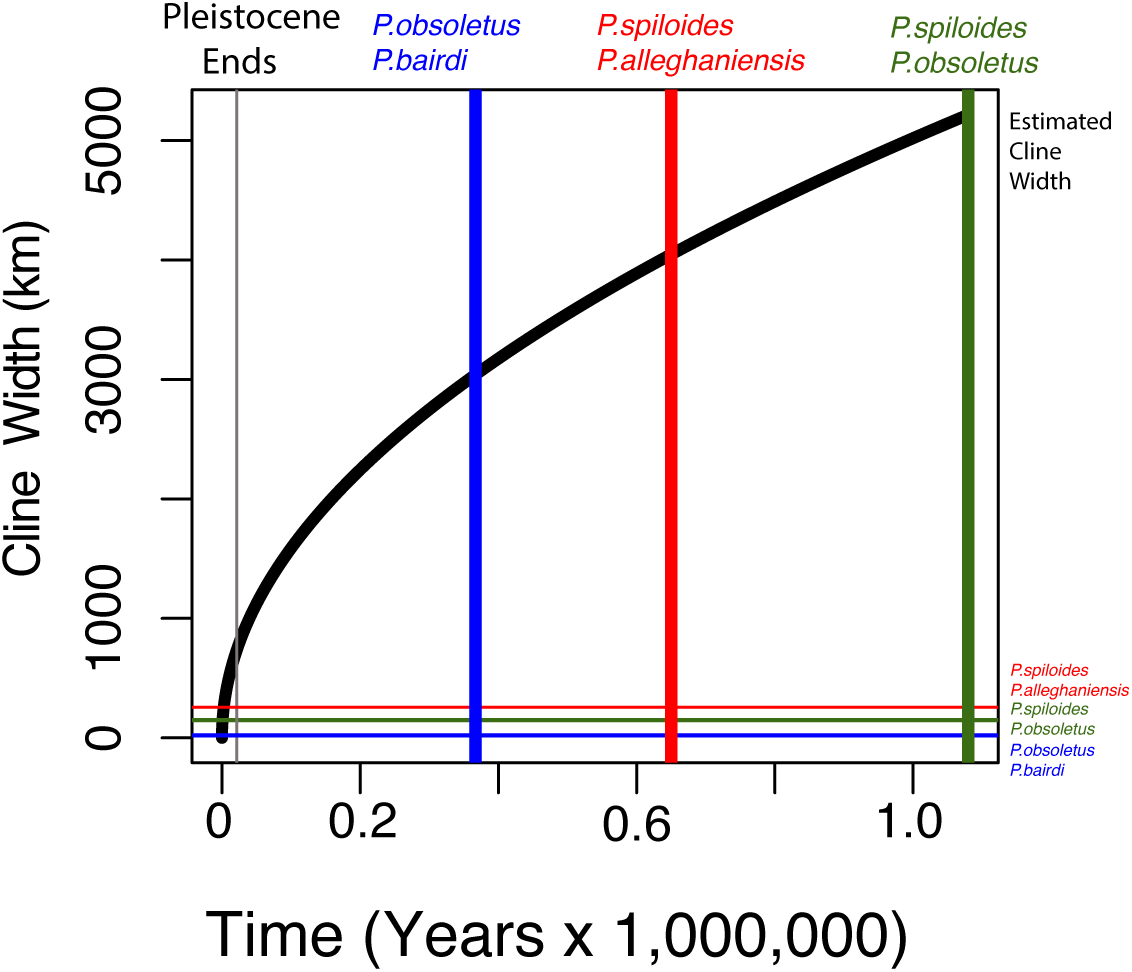
Prediction of cline width under neutrality (black line through time) given divergence times for each species pair: A) *Pantherophis alleghaniensis/P. spiloides*, B) *P. spiloides/P.obsoletus*, and C) *P. obsoletus/P.bairdi.* Colored vertical lines represent times of divergence between species pairs and horizontal colored lines indicate actual cline widths.

**Table S1.**
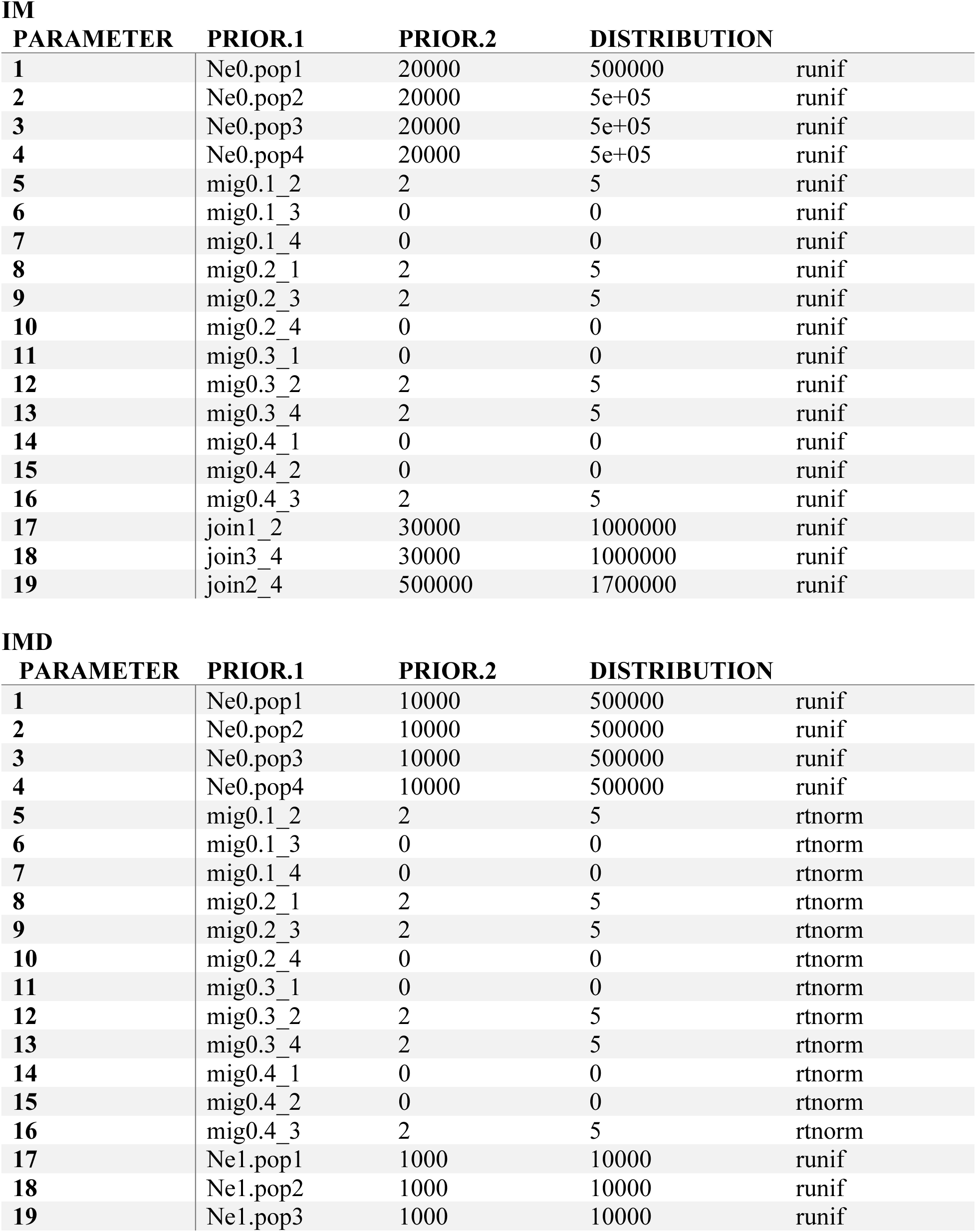
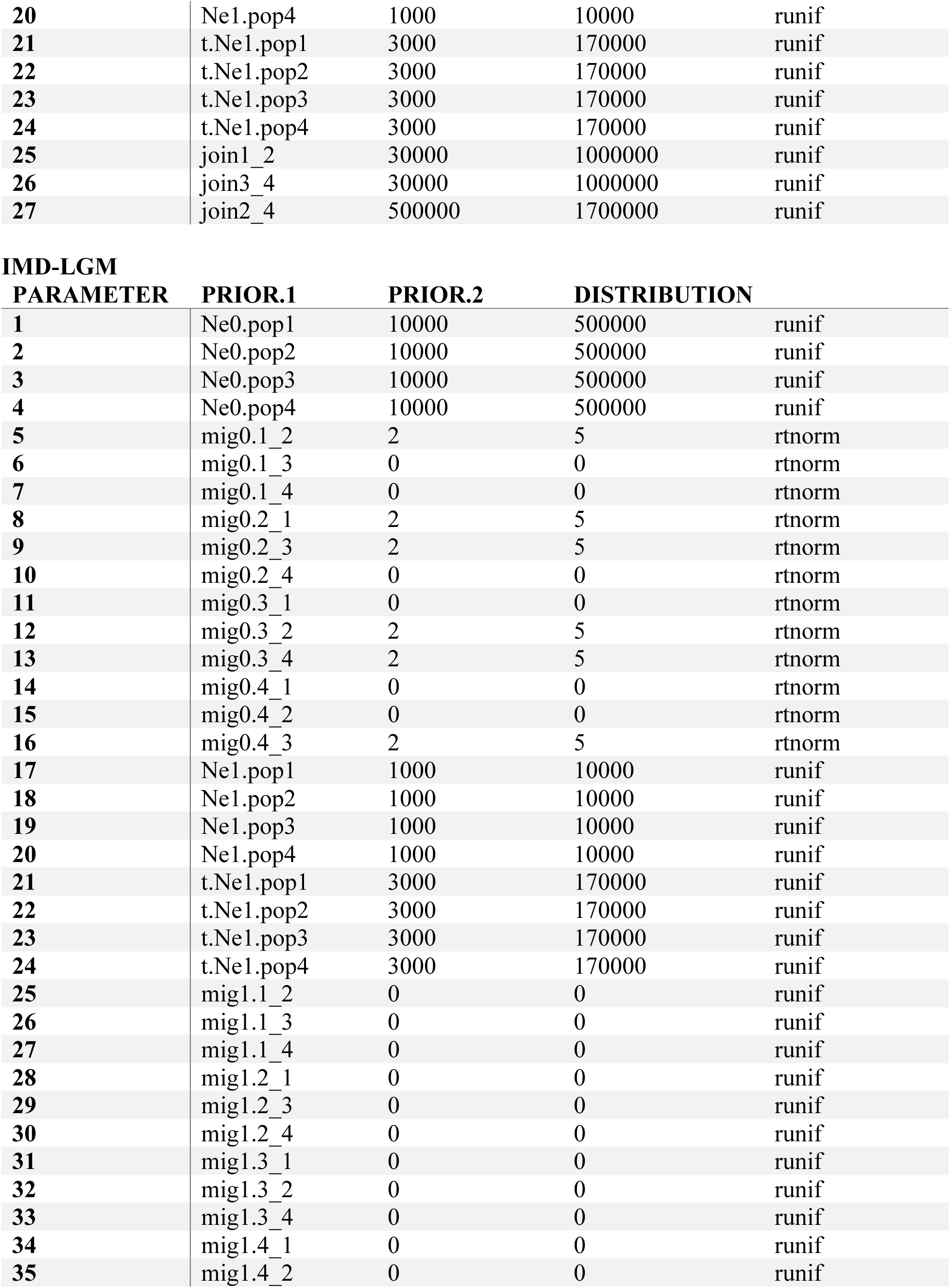
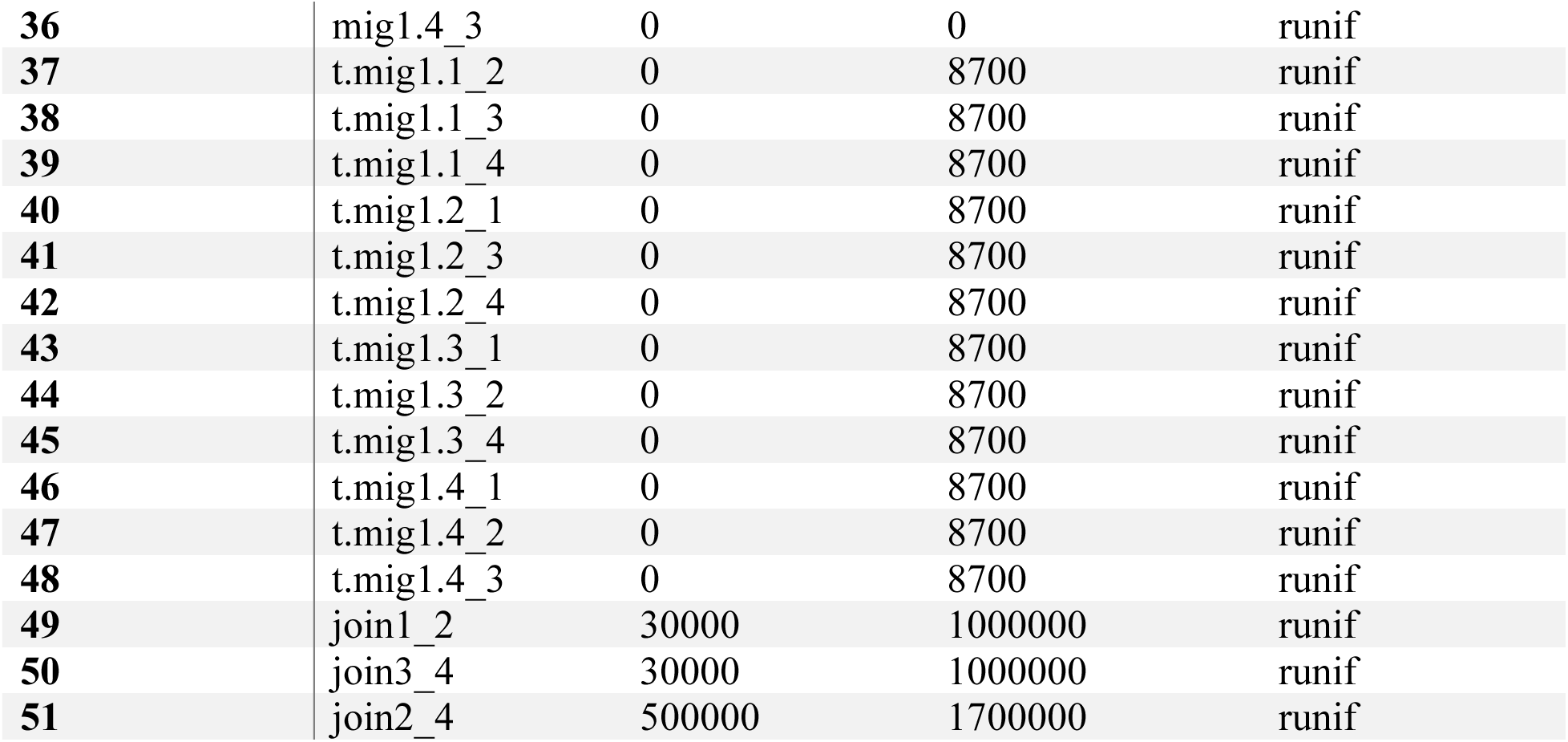
Priors and summary stats for PipeMaster for the IM, IMD, IMD-LGM models; estimating migration, timing of divergence, and historical demography.

| PARAMETER | PRIOR.1 | PRIOR.2 | DISTRIBUTION |  |
| --- | --- | --- | --- | --- |
| 1 | Ne0.pop1 | 20000 | 500000 | runif |
| 2 | Ne0.pop2 | 20000 | 5e+05 | runif |
| 3 | Ne0.pop3 | 20000 | 5e+05 | runif |
| 4 | Ne0.pop4 | 20000 | 5e+05 | runif |
| 5 | mig0.1_2 | 2 | 5 | runif |
| 6 | mig0.1_3 | 0 | 0 | runif |
| 7 | mig0.1_4 | 0 | 0 | runif |
| 8 | mig0.2_1 | 2 | 5 | runif |
| 9 | mig0.2_3 | 2 | 5 | runif |
| 10 | mig0.2_4 | 0 | 0 | runif |
| 11 | mig0.3_1 | 0 | 0 | runif |
| 12 | mig0.3_2 | 2 | 5 | runif |
| 13 | mig0.3_4 | 2 | 5 | runif |
| 14 | mig0.4_1 | 0 | 0 | runif |
| 15 | mig0.4_2 | 0 | 0 | runif |
| 16 | mig0.4_3 | 2 | 5 | runif |
| 17 | join1_2 | 30000 | 1000000 | runif |
| 18 | join3_4 | 30000 | 1000000 | runif |
| 19 | join2_4 | 500000 | 1700000 | runif |

IMD
| PARAMETER | PRIOR.1 | PRIOR.2 | DISTRIBUTION |  |
| --- | --- | --- | --- | --- |
| 1 | Ne0.pop1 | 10000 | 500000 | runif |
| 2 | Ne0.pop2 | 10000 | 500000 | runif |
| 3 | Ne0.pop3 | 10000 | 500000 | runif |
| 4 | Ne0.pop4 | 10000 | 500000 | runif |
| 5 | mig0.1_2 | 2 | 5 | rtnorm |
| 6 | mig0.1_3 | 0 | 0 | rtnorm |
| 7 | mig0.1_4 | 0 | 0 | rtnorm |
| 8 | mig0.2_1 | 2 | 5 | rtnorm |
| 9 | mig0.2_3 | 2 | 5 | rtnorm |
| 10 | mig0.2_4 | 0 | 0 | rtnorm |
| 11 | mig0.3_1 | 0 | 0 | rtnorm |
| 12 | mig0.3_2 | 2 | 5 | rtnorm |
| 13 | mig0.3_4 | 2 | 5 | rtnorm |
| 14 | mig0.4_1 | 0 | 0 | rtnorm |
| 15 | mig0.4_2 | 0 | 0 | rtnorm |
| 16 | mig0.4_3 | 2 | 5 | rtnorm |
| 17 | Ne1.pop1 | 1000 | 10000 | runif |
| 18 | Ne1.pop2 | 1000 | 10000 | runif |
| 19 | Ne1.pop3 | 1000 | 10000 | runif |
| 20 | Ne1.pop4 | 1000 | 10000 | runif |
| 21 | t.Ne1.pop1 | 3000 | 170000 | runif |
| 22 | t.Ne1.pop2 | 3000 | 170000 | runif |
| 23 | t.Ne1.pop3 | 3000 | 170000 | runif |
| 24 | t.Ne1.pop4 | 3000 | 170000 | runif |
| 25 | join1_2 | 30000 | 1000000 | runif |
| 26 | join3_4 | 30000 | 1000000 | runif |
| 27 | join2_4 | 500000 | 1700000 | runif |

IMD-LGM
| PARAMETER | PRIOR.1 | PRIOR.2 | DISTRIBUTION |  |
| --- | --- | --- | --- | --- |
| 1 | Ne0.pop1 | 10000 | 500000 | runif |
| 2 | Ne0.pop2 | 10000 | 500000 | runif |
| 3 | Ne0.pop3 | 10000 | 500000 | runif |
| 4 | Ne0.pop4 | 10000 | 500000 | runif |
| 5 | mig0.1_2 | 2 | 5 | rtnorm |
| 6 | mig0.1_3 | 0 | 0 | rtnorm |
| 7 | mig0.1_4 | 0 | 0 | rtnorm |
| 8 | mig0.2_1 | 2 | 5 | rtnorm |
| 9 | mig0.2_3 | 2 | 5 | rtnorm |
| 10 | mig0.2_4 | 0 | 0 | rtnorm |
| 11 | mig0.3_1 | 0 | 0 | rtnorm |
| 12 | mig0.3_2 | 2 | 5 | rtnorm |
| 13 | mig0.3_4 | 2 | 5 | rtnorm |
| 14 | mig0.4_1 | 0 | 0 | rtnorm |
| 15 | mig0.4_2 | 0 | 0 | rtnorm |
| 16 | mig0.4_3 | 2 | 5 | rtnorm |
| 17 | Ne1.pop1 | 1000 | 10000 | runif |
| 18 | Ne1.pop2 | 1000 | 10000 | runif |
| 19 | Ne1.pop3 | 1000 | 10000 | runif |
| 20 | Ne1.pop4 | 1000 | 10000 | runif |
| 21 | t.Ne1.pop1 | 3000 | 170000 | runif |
| 22 | t.Ne1.pop2 | 3000 | 170000 | runif |
| 23 | t.Ne1.pop3 | 3000 | 170000 | runif |
| 24 | t.Ne1.pop4 | 3000 | 170000 | runif |
| 25 | mig1.1_2 | 0 | 0 | runif |
| 26 | mig1.1_3 | 0 | 0 | runif |
| 27 | mig1.1_4 | 0 | 0 | runif |
| 28 | mig1.2_1 | 0 | 0 | runif |
| 29 | mig1.2_3 | 0 | 0 | runif |
| 30 | mig1.2_4 | 0 | 0 | runif |
| 31 | mig1.3_1 | 0 | 0 | runif |
| 32 | mig1.3_2 | 0 | 0 | runif |
| 33 | mig1.3_4 | 0 | 0 | runif |
| 34 | mig1.4_1 | 0 | 0 | runif |
| 35 | mig1.4_2 | 0 | 0 | runif |
| 36 | mig1.4_3 | 0 | 0 | runif |
| 37 | t.mig1.1_2 | 0 | 8700 | runif |
| 38 | t.mig1.1_3 | 0 | 8700 | runif |
| 39 | t.mig1.1_4 | 0 | 8700 | runif |
| 40 | t.mig1.2_1 | 0 | 8700 | runif |
| 41 | t.mig1.2_3 | 0 | 8700 | runif |
| 42 | t.mig1.2_4 | 0 | 8700 | runif |
| 43 | t.mig1.3_1 | 0 | 8700 | runif |
| 44 | t.mig1.3_2 | 0 | 8700 | runif |
| 45 | t.mig1.3_4 | 0 | 8700 | runif |
| 46 | t.mig1.4_1 | 0 | 8700 | runif |
| 47 | t.mig1.4_2 | 0 | 8700 | runif |
| 48 | t.mig1.4_3 | 0 | 8700 | runif |
| 49 | join1_2 | 30000 | 1000000 | runif |
| 50 | join3_4 | 30000 | 1000000 | runif |
| 51 | join2_4 | 500000 | 1700000 | runif |

